# A simple approximation to bias in the genetic effect estimates when multiple disease states share a clinical diagnosis

**DOI:** 10.1101/483198

**Authors:** Iryna Lobach, Inyoung Kim, Alexander Alekseyenko, Siarhei Lobach, Li Zhang

## Abstract

Case-control genome-wide association (CC-GWAS) studies might provide valuable clues to the underlying pathophysiologic mechanisms of complex diseases, such as neurodegenerative disease, cancer. A commonly overlooked complication is that multiple distinct disease states might present with the same set of symptoms and hence share a clinical diagnosis. These disease states can only be distinguished in a biomarker evaluation that might not be feasible on the whole set of cases in the large number of samples that are typically needed for CC-GWAS. Instead, the biomarkers are measured on a subset of cases. Or an external reliability study estimates frequencies of the disease states of interest within the clinically diagnosed set of cases. These frequencies often vary by the genetic and/or non-genetic variables. We derive a simple approximation that relates the genetic effect estimates obtained in a logistic regression model with the clinical diagnosis as an outcome variable to the estimates in the relationship to the true disease state of interest. We performed simulation studies to assess accuracy of the approximation that we’ve derived. We next applied the derived approximation to the analysis of the genetic basis of innate immune system of Alzheimer’s disease.

## INTRODUCTION

Case-control genome-wide analyses scan (CC-GWAS) is a tool that is widely used to elucidate the genetic basis of complex diseases. A common complication is that multiple distinct disease states share the observed symptoms and hence the clinical diagnosis. Frequencies of the disease states within the clinical diagnosis often vary by the key variables. If the disease states have distinct genetic basses, the analyses with a clinical diagnosis as an outcome variable might be substantially biased (Carroll et al, 2006).

The specific example that motivated this study is the analyses of the genetic susceptibility to Alzheimer’s disease (AD). The clinical diagnosis of AD is typically made based on a set of descriptive criteria and only a small subset of cases receives positron emission tomography (PET) to evaluate for amyloid positivity, what is a requirement for the *true*, or pathologically defined, AD. Recent biomarker studies (Salloway and Sperling, 2015) estimate that 36% of ApoE *ε*4 non-carriers and 6% of ApoE *ε*4 carriers diagnosed with AD do not have evidence for amyloid as measured by PET, hence do not qualify for the *true* AD diagnosis.

We are interested to examine the role of the genetic variants serving the innate immune system in susceptibility to AD, i.e. the AD symptoms underlined by the amyloid deposition. The usual analyses define the outcome variable in a regression analysis to be the clinical diagnosis. We, however, recognize heterogeneity of the clinical diagnosis where the underlying disease state separates the cases into a subset with amyloid-related AD, what is the disease state of interest; and non-amyloid-related AD, what is the nuisance disease state. We derive the theoretical approximation that provides a simple and general relationship between *B* and Γ estimates using Kullback-Leibler divergence (Kullback, 1959).

Our paper is organized as follows. First, in the Material and Methods section we present the setting, notation, and the proposed approximation for various models. Next, in the Simulation Experiments section we describe the empirical studies that are conducted to compare the resulting performance of the approximation that we derived relative to the average observed across many simulated datasets. We then compare the estimates in a practical setting of an Alzheimer’s disease study that aims to investigate the genetic basis of innate immune system in the relationship to the AD symptoms underlined by amyloid pathology. We conclude our paper with brief Discussion.

## MATERIALS AND METHODS

We define *G* to be the genotype of single nucleotide polymorphisms (SNPs) measured at multiple locations. Let *X* and *Z* be the environmental variables that might interact. We assume that the genotype is independent of the environment and follows Hardy-Weinberg equilibrium model *Q* (*g*; *θ*), where *θ* is the frequency of the minor allele.

We define *D*^*CL*^ be the observed clinical diagnosis that is inferred based on a set of descriptive criteria that characterize symptoms. Let *D* denote the true disease states, where *D* = 1 indicates the disease state of interest and *D* = 1^*^is the nuisance disease state. It might not be possible to measure *D* on the set of cases in a GWAS, instead *D* is available on a subset or frequencies of *D* within the clinically defined set of cases are reliably estimated in an external reliability study. We define the clinical-pathological diagnosis relationship using *τ* (*X*) = *pr*(*D*= 1|*D*^*CL*^= 1,*X*), what is a frequency of the disease state of interest within the clinically diagnosed set and the frequency varies by *X*. In the context of AD study, *pr* (*D*= 1^*^|*D*^*CL*^= 1,*X*) =1−*π*4; (*X*), *pr*(*D*= 0|*D*^*CL*^= 1,*X*) = 0, *pr*(*D*= 1^*^|*D*^*CL*^= 0,*X*) = *pr*(*D*= 1|*D*^*CL*^= 0,*X*) =0 and *pr*(*D*= 1^*^|*D*^*CL*^= 0,*X*) = *pr*(*D*= 0|*D*^*CL*^= 0,*X*) = 1. We define the probabilities of the clinical diagnosis in the population to be *π*_*d*_*CL* = *pr*(*D*^*CL*^ = *d*^*cl*^). Similarly, we let frequencies of the true pathologic state in the population to be *π*_d_= *pr*(*D* = *d*).

For clarity of presentation we assume that genotype is binary to indicate presence of a minor allele, environmental variables *X* and *Z* are Bernoulli with frequencies *η*_*X*_and *η*_*Z*_, respectively. In the Appendix we discuss how to extend the approximation to the categorical and continuous variables.

### Model 1. *β*_*G*_

We first consider a setting when only the genetic variable *G* is in the risk model, i.e. the true disease risk model is

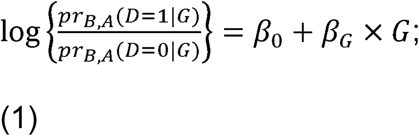

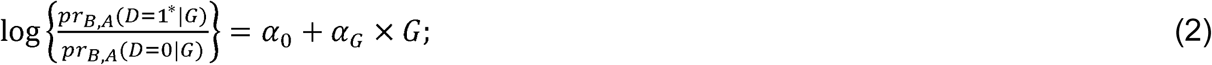

while the model used is the usual logistic regression model with the clinical diagnosis as an outcome variable, i.e.

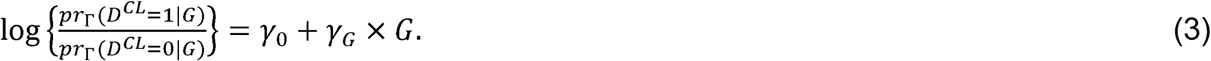

Derivations provided in Appendix A1 show that

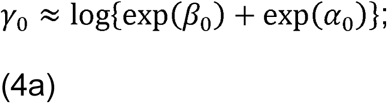

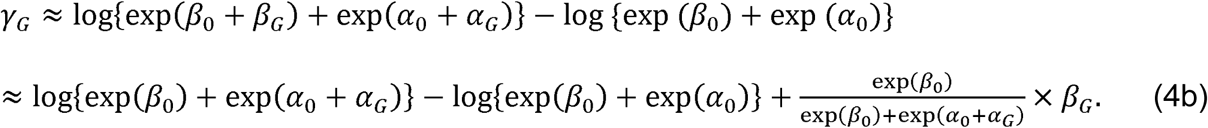

From (4a) and (4b), we derive that

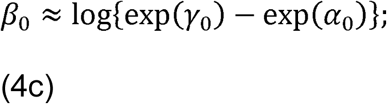

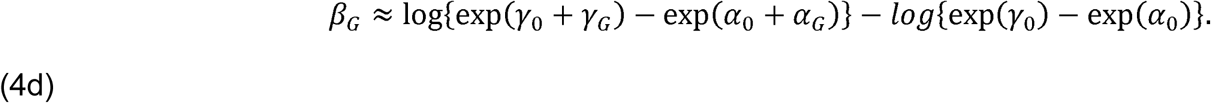

Appendix A3 describes how to obtain *β*_0_, *β*_*G*_, *α*_0_, *α*_*G*_, assuming estimates of *γ*_0_, *γ*_*G*_ are available from the usual logistic regression and reliable estimates of *τ* = *τ* (1) × *pr*(*X* = 1) + *τ* (0) × *pr*(*X* = 0) and *π*_1_are available in the literature.

### Model 2. *β*_*G*_ and *β*_*X*_

We next consider a setting when the genetic variable *G* and an environmental variable *X* are in the risk model, i.e. the true disease risk model is

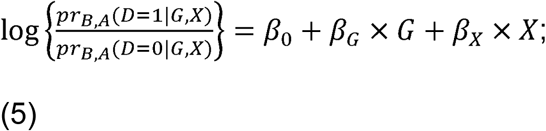

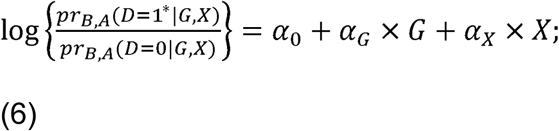

while the model used is

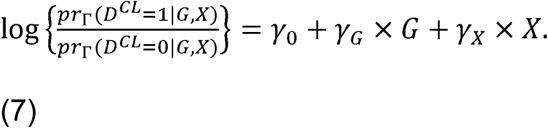

Derivations provided in Appendix A2 show that

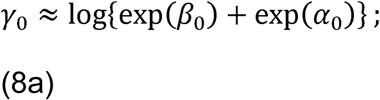

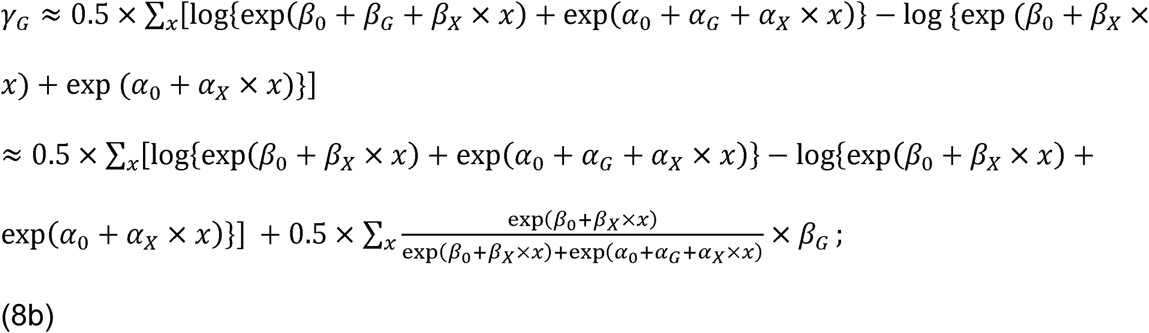

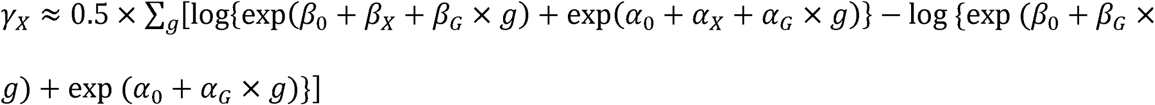

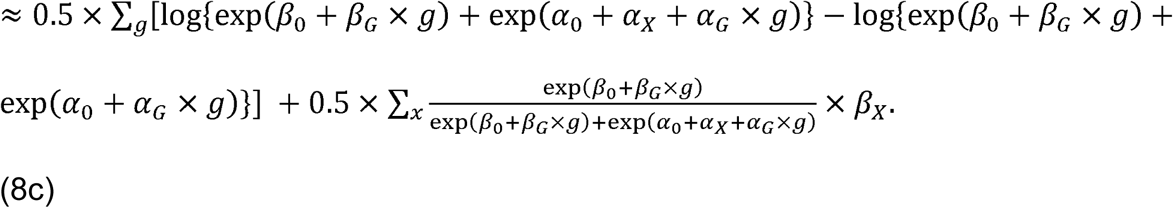

### Model 3

***β*_*G*_, *β*_*X*_, *β*_*Z*_, and *β*_*X*×*Z*.:_** A model with interaction between the environmental variables *X* and *Z* is discussed in Appendix.

### Model 4

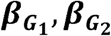 **and** 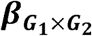: A model with gene-gene interactions is discussed in Appendix.

### Remarks

1. Model 1, equation (4b). If *β*_*G*_= *α*_*G*_= 0, then *γ*_*G*_= 0.
2. Model 2, equation (8b). If *β*_*G*_= *α*_*G*_= 0, then *γ*_*G*_= 0.
3. Model 2, equation (8c). If *β*_*X*_= *α*_*X*_= 0, then *γ*_*X*_= 0.
4. Remarks 1-3 describe when the usual logistic regression models with the clinical diagnosis as an outcome variable correctly estimate the null effect.
5. The equations that we derived apply to several possible likelihood functions. For example, parameter estimates in Model 3 can be estimated based on the usual logistic regression model, i.e. the probability of the form *pr* _Γ_(*D*^*CL*^|*G,X,Z*) or in a pseudolikelihood (Spinka et al, 2005; Lobach et al, 2018) *pr*_Γ_(*D*^*CL*^,*G*|*X,Z, β*4; = 1), where, *β*4; = 1 is an imaginary indicator of being selected into the study. All the derivations apply to both models.

## SIMULATION STUDIES

### False positive rate

We first perform a series of simulation experiments to examine a false positive rate in the estimates of *β*_*G*_ when the data are simulated from model (1)-(2), but the parameter estimates are obtained from model (3). We define the false positive rate to be the fraction of p-values≤0.05 across 10,000 simulated datasets in the usual logistic regression analyses as an outcome variable, i.e. (3), when in fact *β*_*G*_= 0. We simulate the data using model (1) with coefficients *β*_0_= 0.5,*β*_*G*_= 0,*α*_*G*_= log(1) = 0,log(1.5) = 0.41,log (2) = 0.69. We next estimate parameters using model (3). **Table 1** presents false positive rates in datasets with *n*_0_ = *n*_1_= 3,000;10,000. When the genetic effect is not associated with the clinical diagnosis, the false positive rate is nominal, i.e. is nearly 0.05. When *α*_*G*_increases, the false positive rate gets inflated, e.g. when *α*_*G*_= log(1.5) = 0.41, the false positive rate is 0.72. Increase in sample size did not result in decrease of the false positive rate.

**Table 1:** False positive rate defined as the proportion of p-values0.05 across 10,000 simulated datasets in the usual logistic regression analyses as an outcome variable (3), when in fact *β*_*G*_=0 and the data are generated from (1)-(2). We let *β*_0_ = 0.5, *β*_*G*_ = 0, *α*_*G*_ = log(1) = 0,log(1.5) = 0.41,log (2) = 0.69.

| $\alpha_G =$ | $\text{Log}(1) = 0$ | $\text{Log}(1.5) = 0.41$ | $\text{Log}(2) = 0.69$ |
| --- | --- | --- | --- |
| $n_0 = n_1 = 3,000$ | 0.052 | 0.72 | 0.99 |
| $n_0 = n_1 = 10,000$ | 0.048 | 0.79 | 0.99 |

## Approximation vs. empirical estimates

We next perform a series of simulation experiments to assess the magnitude of bias and the approximation to the relationships that we’ve derived. First, we estimate the bias empirically as the average across 500 simulated datasets where the data are simulated using the true model (1)-(2), (5)-(6), (A3)-(A4) based on coefficients *B* and A, but estimate the parameters Γ in the usual logistic regression model (3), (7) and (A5). We then compare these averages to the approximations that we’ve derived.

We simulate genotype (*G*), age (*A*), sex (*S*), ApoE *ϵ*4 status to be Bernoulli with frequencies *θ*_*G*_,*θ*_*A*_,*θ*_*S*_,*θ*_*ϵ*4_. In the context of previous notations, *X* is the ApoE *ϵ*4 status and *Z* is a set consisting of *G,A,S*. We then simulated the clinical diagnosis status *D*^*CL*^according to the models (3), (7) and (A5) and the true disease states *D* according to model (1)-(2), (5)-(6), (A3)-(A4). In all simulations we let *θ*_*G*_ = 0.10, *θ*_*A*_ = 0.50, *θ*_*S*_ = 0.52, *θ*_*ϵ*4_ = 0.07.

### Model 1

We fist simulate the data using model (1)-(2) and estimate parameters in the logistic model (3). We set *β*_0_ = 0.5, *β*_*G*_ = log(1) = 0,log(1.5) = 0.41,log(2) = 0.69,log(2.5) = 0.92, log(3)=1.1, *α*_*G*_ = log(1) = 0,log(1.5) = 0.41,= 0.69 and simulate datasets with 3,000 cases and 3,000 controls. **Table 2** presents empirical estimates of *β*_*G*_ and the approximation (4b). Across all values of *β*_*G*_and *α*_*G*_, the approximation (4b) is accurate relative to the empirical estimate.

**Table 2:** Empirical estimates of *β*_*G*_and *approximation* (4b). The data are simulated from models (1)-(2) and is estimated using model (3). Empirical estimates are the averages across 500 datasets with 3,000 cases and 3,000 controls. We let *β*_0_ = *α*_0_ = 0.5, *β*_*G*_ = log(1) = 0,log(1.5) = 0.41,log(2) = 0.69,log(2.5) = 0.92, log(3) = 1.1, *β*_*ϵ*4_ = *α*_*ϵ*4_ = log(8), *α*_*G*_ = log(1) = 0,log(2) = 0.41,log(3) = 0.69,log(4) = 1.1.

| $\alpha_G$ | $\beta_G$ | | | | |
| --- | --- | --- | --- | --- | --- |
|  | Log(1)=0 | Log(1.5)=0.41 | Log(2)=0.69 | Log(2.5)=0.92 | Log(3)=1.1 |
| Log(1)=0 | 0.003, 0 | 0.23, 0.22 | 0.41, 0.40 | 0.57, 0.56 | 0.70, 0.69 |
| Log(2)=0.41 | 0.41, 0.41 | 0.57, 0.56 | 0.70, 0.69 | 0.82, 0.81 | 0.92, 0.92 |
| Log(3)=0.69 | 0.70, 0.69 | 0.82, 0.81 | 0.93, 0.92 | 1.0, 1.0 | 1.1, 1.1 |
| Log(4)=1.1 | 0.93, 0.92 | 1.02, 1.01 | 1.1, 1.1 | 1.2, 1.2 | 1.3, 1.3 |

### Model 2

We next generate data using models (5)-(6) but estimate parameters using model (7). We let *β*_0_ = *α*_0_ = 0.5, *β*_*G*_ = log(1) = 0,log(1.5) = 0.41,log(2) = 0.69,log(2.5) = 0.92,log(3) = 1.1,*β*_*ϵ*4_ = *α*_*ϵ*4_ = log(8),*α*_*G*_ = log(1) = 0,log(2) = 0.41,log(3) = 0.69,log(4) = 1.1 and generate datasets with 3,000 cases and 3,000 controls. Approximations and the empirical estimates for *γ*_G_ shown in **Table 3** demonstrate that the approximation (8b) is accurate relative to the empirical estimates. The empirical estimate of *γ*_*ϵ*4_is 2.09, while the approximation is 2.08.

**Table 3:** Empirical estimates of *γ*_*G*_and *approximation* (8b). The data are simulated from models (5)-(6) and is estimated using model (7). Empirical estimates are the averages across 500 datasets with 3,000 cases and 3,000 controls. We let *β*_0_ = *α*_0_ = 0.5, *β*_*G*_ = log(1),log(1.5), log(2), log(2.5), log(3),*β*_*ϵ*4_ = *α*_*ϵ*4_ = log(8), *α*_*G*_ = log(1), log(2), log(3), log(4).

| $\alpha_G$ | $\beta_G$ | | | | |
| --- | --- | --- | --- | --- | --- |
|  | Log(1)=0 | Log(1.5)=0.41 | Log(2)=0.69 | Log(2.5)=0.92 | Log(3)=1.1 |
| Log(1)=0 | 0.0056, 0 | 0.23, 0.22 | 0.41, 0.40 | 0.57, 0.56 | 0.70, 0.69 |
| Log(2)=0.41 | 0.41, 0.41 | 0.57, 0.56 | 0.79, 0.69 | 0.82, 0.81 | 0.92, 0.92 |
| Log(3)=0.69 | 0.70, 0.69 | 0.82, 0.81 | 0.92, 0.92 | 1.0, 1.0 | 1.1, 1.1 |
| Log(4)=1.1 | 0.92, 0.92 | 1.02, 1.01 | 1.1, 1.1 | 1.2, 1.2 | 1.2, 1.3 |

### Model 3

We next simulate data using models (A3)-(A4) and estimate parameters using model (A5), with the approximation derived in (A6a-c).

### Setting 1

We first consider a setting when the nuisance disease is not associated with the genotype (*α*_*G*_ = 0) and when *ϵ*4 and A × *ϵ*4 are not associated with the nuisance disease status (*α*_*ϵ*4_ = 0,*α*_*G*_ = 0,*α*_*A*×*ϵ*4_ = 0). We simulate the clinical diagnosis and disease states with coefficients *β*_0_ = −1,*β*_*S*_ = *log*(0.92) = −0.08,*β*_*ϵ*4_ = *log*(8) = 2.1,*β*_*S*_ = *log*(2) = 0.69, *β*_*G*_ = *log*(1),*log*(1.5),*log*(2),*log*(2.5),*log*(3),*β*_*A*×*ϵ*4_ = *log*(1),*log*(2),*log*(3),*log*(3), *α*_0_ = −1,*α*_*S*_ = *log*(0.92),*α*_*ϵ*4_ = 0,*α*_*A*_ = *log*(2),*α*_*G*_ = 0,*α*_*A*×*ϵ*4_ = 0.

**Figure 1** presents biases in estimates of *β*_*G*_(panel A), *β*_*A*_(panel B), *β*_*S*_(panel C), *β*_*ϵ*4_(panel D) and *β*_*A×ϵ*4_(panel E) in studies with 3,000 cases and 3,000 controls; values of *β*_*A ×ϵ*4_are shown along the x-axis and values of *β*_*G*_are indicated by color. **Figure 1** panels **A** and **D** show that bias in the estimates of *β*_*G*_and *β*_*ϵ*4_can be substantial with largest bias of −0.06; panel **E** shows that bias in *β*_*A×ϵ*4_is notable in this case ranging from 0.01 to −0.06; estimates of *β*_*A*_and *β*_*S*_are nearly unbiased consistent with the theoretical observations that the null effect in some settings can be estimated with no bias even in a misspecified model. We note that magnitude of bias in 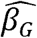 and 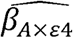 increases as the true value of the coefficient increases.

Shown on **Figure 2** are the empirical bias (Emp) and the approximation (AX) of bias in *β*_*G*_indicated by color with values of *β*_*A* ×*ϵ*4_along the x-axis and values of *β*_*G*_along the panels. The difference between the Emp and AX starts at ≈ 0.6 when *β*_*G*_ = 0.41 and increases to ≈ 1.2 when *β*_*G*_ = 1.1. Bias of 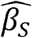 and 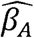 is approximated to be <0.0001. Shown on **Figure 3** are Emp and AX of estimates of *β*_*ϵ*4_, and **Figure 4** is presenting estimates of *β*_*A*×*ϵ*4_.

### Setting 2

We next simulate datasets with 30,000 cases and 30,000 controls in the Setting 1. **Supplementary Figure 1** shows that biases in the estimates noted in Setting 1 persists for larger sample sizes.

### Setting 3

We next consider a setting when with the nuisance disease the genotype is associated (*α*_*G*_ = *log*(1.5)), *α*_*ϵ*4_is associated (*α*_*ϵ*4_ = *log*(2)), and no interaction *α*_*A*×*ϵ*4_ = 0. We next change the parameters for the nuisance state to be *α*_0_ = −1,*α*_*S*_ = *log*(0.92),*α*_*ϵ*4_ = *log*(2),*α*_A_ = *log*(2),*α*_*G*_ = *log*(1.5),*α*_*A*×*ϵ*4_ =0 and all other parameters as in Setting 1. Shown on **Supplementary Figure 2** are biases in the estimates of the parameters of interest that reach −1.4 for 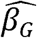, are near −0.5 for 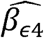, and can reach −0.15 for 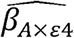.

### Setting 4

We next consider a Setting 1 but with more common disease of interest, i.e. *β*_0_ = 1.5. **Supplementary Figure 3** is showing empirical bias in all estimates. The estimates can still be substantially biased.

### Setting 5

We next consider a setting where the genetic variable is associated with the nuisance disease state (*α*_*G*_ = *log*(2)) and there is significant *A* ×*ϵ*4 interaction (*α*_*A*×*ϵ*4_ = *log*(2)). We next change the parameters for the nuisance state to be *α*_0_ = 0.5,*α*_*S*_ = *log*(0.80),*α*_*ϵ*4_ = *log*(4),*α*_*A*_ = *log*(3),*α*_*G*_ = *log*(2),*α*_*A*×*ϵ*4_ = *log*(2). **Figure 5** presents biases in the estimates and **Supplementary Figures 4-5** show the empirical estimates and the approximations.

**Supplementary Figure 7:** Frequency of the disease state of interest (*D* = 1) and the nuisance disease (*D* = 1^*^) when *β*_0_ = 1.5,*β*_*S*_ = *log*(0.80),*β*_*ϵ*4_ = *log*(8),*β*_*A*_ = *log*(3), *β*_*G*_ = *log*(1),*log*(1.5),*log*(2),*log*(2.5),*log*(3),*β*_*G* ×*ϵ*4_ = *log*(1),*log*(2),*log*(3),*log*(3), *α*_0_ = 0.5,*α*_*S*_ = *log*(0.80),*α*_*ϵ*4_ = *log*(4),*α*_*A*_ = *log*(3),*α*_*G*_ = *log*(2),*α*_*A*×*ϵ*4_ = *log*(2), *θ*_*G*_ = 0.10, *θ*_*A*_ = 0.50, *θ*_*S*_ = 0.52, *θ*_*ϵ*4_ = 0.07. Shown along the x-axis are values of *β*_*G*_and indicated by color are values of *β*_*A*×*ϵ*4_. We note that these frequencies are similar to those in context of Alzheimer’s disease.

## ROLE OF THE GENETIC VARIANTS SERVING INNATE IMMUNE SYSTEM IN SUSCEPTIBILITY TO ALZHEIMER’s DISEASE

We apply the usual logistic analyses with the clinical diagnosis as an outcome variable to a dataset collected as part of the Alzheimer’s Disease Genetics Consortium. We next apply the approximations (7)-(10) and (11)-(14) to see how the genetic estimates change when presence of the nuisance disease state is recognized.

We mapped Illumina Human 660K markers onto human chromosomes using NCBI dbSNP database (https://www.ncbi.nlm.nih.gov/projects/SNP/). Chromosomal location, proximal gene or genes and gene structure location (e.g. intron, exon, intergenic, UTR) has been recorded for all SNPs. From these data we inferred 165 SNPs to reside in genes serving innate immune system.

The dataset consists of 727 controls and 2,797 cases diagnosed with AD.

We are interested to examine a relationship between the pathologic disease state of AD characterized by presence of amyloid deposition and each of the 165 SNPs serving the innate immune system. We include ApoE *ϵ*4 status, age, and sex in the model with an interaction between ApoE and age. The genetic variant is modeled as a Bernoulli variable as an indicator of presence or absence of a minor allele. Age is Bernoulli as well that corresponds to a median split in the dataset.

**Table 4** presents estimates of effects of the SNPs obtained using the usual logistic regression model with the clinical diagnosis as an outcome variable in a univariable model (3) and with adjustment for SNP + ApoE ε4 + Age + Sex (7); and the corresponding models (1-2) and (5-6) that recognize presence of the nuisance disease state. In the univariable setting the empirical bias is estimated as the difference between the main effect estimates obtained in model (3) and model (1-2), and the approximation to the bias is estimated as derived in (4b). In the multivariable setting, the empirical bias is the difference between main effect estimates obtained in model (7) and (5-6), and the approximation is as derived in (8b).

**Table 4:**
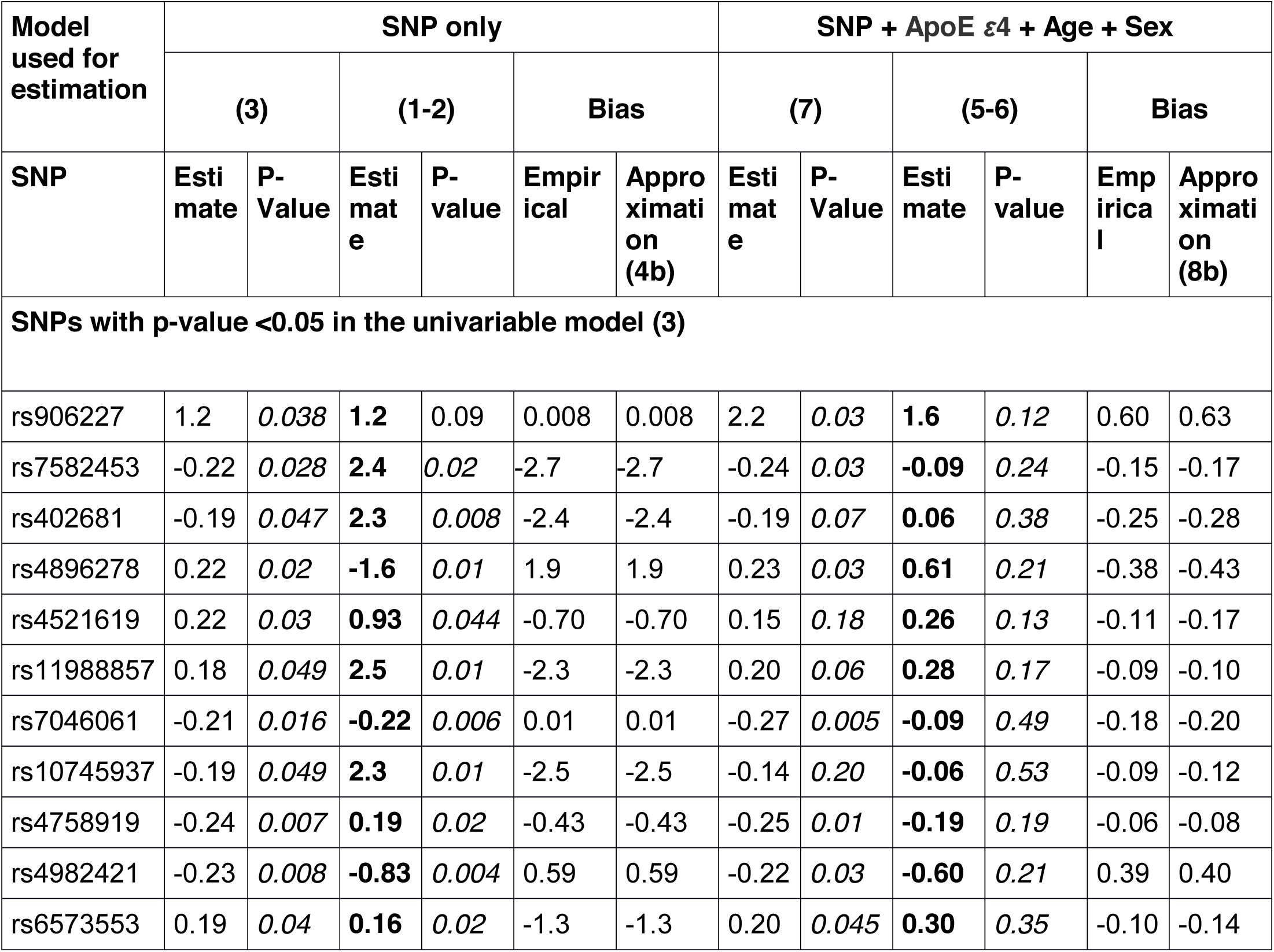
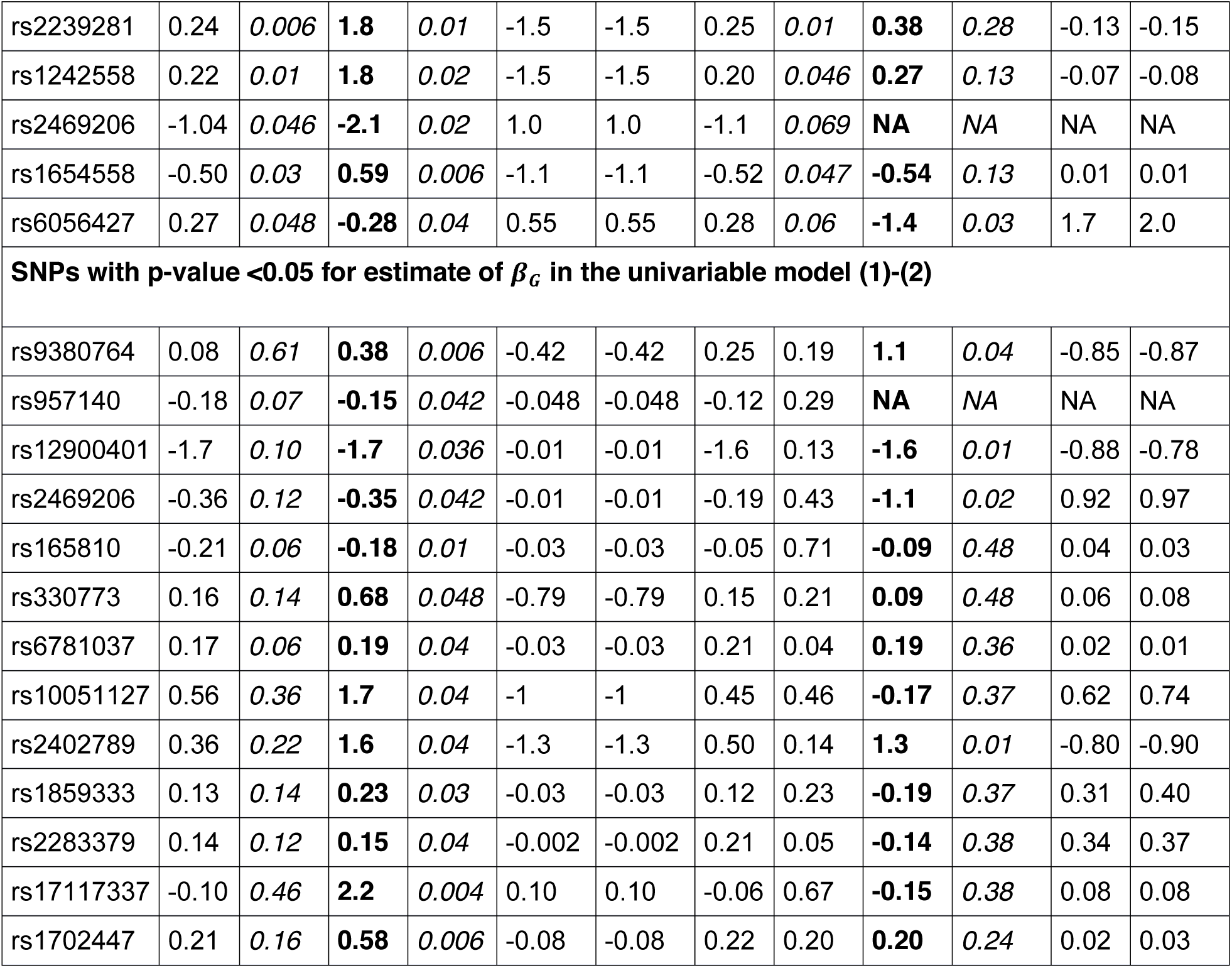
Main effect estimates of SNPs obtained using the usual logistic regression with the clinical diagnosis as an outcome variable in a univariable model (3) and with adjustment for SNP + ApoE ε4 + Age + Sex (7); and the corresponding models (1-2) and (5-6) that recognize presence of the nuisance disease state. In the univariable setting the empirical bias is estimated as the difference between the main effect estimates obtained in model (3) and model (1-2), and the approximation is as derived in (4b). In the multivariable model, the empirical bias is the difference between main effect estimates obtained in model (7) and (5-6), and the approximation is as derived in (8b).

First shown in **Table 4** are 16 estimates with p-value<0.05 after the Benjamini-Hochberg multiple testing adjustment in a univariable model (3) and then added are 13 SNPs with p-value <0.05 in a univariable model (1-2). Across all these SNPs, the approximation was accurate relative to the empirical bias.

## DISCUSSION

We’ve examined a situation when multiple disease states share observed symptoms and hence the clinical diagnosis. Both theoretically and in extensive simulation studies we observed that the magnitude of bias can be substantial in situations when frequency of the nuisance disease state within the clinically diagnosed set varies by the key variables. We derived a simple and general approximation to the relationship between the genetic effect estimates that use the clinical diagnosis as an outcome variable and the estimates that recognize presence of the nuisance disease state.

While the effect of misclassification of the disease status has been examined extensively in statistical literature (Carroll et al, 2006), we extend the literature by deriving a simple and general approximation to the bias in a multivariable setting. The approximation provides a simple formula to assess how elastic the estimates of interest are to the values of parameters in the nuisance risk model. The regression coefficients or plausible ranges for the coefficients of the nuisance disease state are often available in the literature.

Simulation studies that we conducted showed that when presence of the nuisance disease is ignored, the genetic effect estimates can be biased in either direction. These biases can be substantial in magnitude leading to false positive and false negative results.

While our study is motivated by the setting of Alzheimer’s disease, the results are readily applicable for other complex diseases. For example, Manchia el al (2013) examined the effect of heterogeneity, i.e. presence of non-cases, in the context of diabetes and showed that ignoring the heterogeneity leads to reduced statistical power to detect an association and also reduced the estimated risks attributable to susceptibility alleles.

The approximation that we’ve derived is widely applicable in other areas of research where the diagnosis is heterogeneous. For example, when disease states correspond to subtypes of a complex disease. We also see the application to the analyses of Electronic Health Records, where the disease status might be subject to exposure-dependent differential misclassification (Chen et al, 2017).

## Supporting information

## ACKNOWLEDGEMENTS

Dr. Lobach is supported by 5R21AG043710-02.

Genotyping is performed by Alzheimer’s Disease Genetics Consortium (ADGC), U01 AG032984, RC2 AG036528. Phenotypic collection is coordinated by the National Alzheimer’s Coordinating Center (NACC), U01 AG016976

Samples from the National Cell Repository for Alzheimer’s Disease (NCRAD), which receives government support under a cooperative agreement grant (U24 AG21886) awarded by the National Institute on Aging (NIA), were used in this study. We thank contributors who collected samples used in this study, as well as patients and their families, whose help and participation made this work possible;

Data for this study were prepared, archived, and distributed by the National Institute on Aging Alzheimer’s Disease Data Storage Site (NIAGADS) at the University of Pennsylvania (U24-AG041689-01)

We thank Ivan Belousov for help with the computations.

## APPENDIX

### A1. Approximation using Kullback-Leibler divergence

We show schematics of the derivations based on Model 3, the other models can be derived accordingly. We denote the model the true model (9)-(10) based on probability *pr*_Γ_(*D*^*CL*^|*G,X,Z*) or *pr*_Γ_(*D*^*CL*^,*G*|*X,Z, δ* = 1) as *Q*_*B,A*_(*D*^*CL*^, *G,X,Z*) = *pr*_*B,A*_(*D*^*CL*^,*G*|*X, Z, δ* = 1). Similarly, we denote model (3) with (4) as *Q*_Γ_(*D*^*CL*^,*G,X,Z*) = *pr*_Γ_ (*D*^*CL*^| *G,X,Z*). Kullback (1959) showed that parameters Γ converge to values that minimize Kullback-Leibler divergence criteria between the two models, specifically

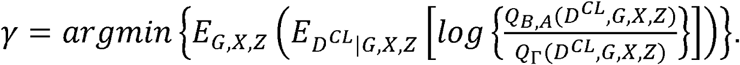

Considerable algebraic derivations arrive to the following system of equations to be solved for parameters Γ

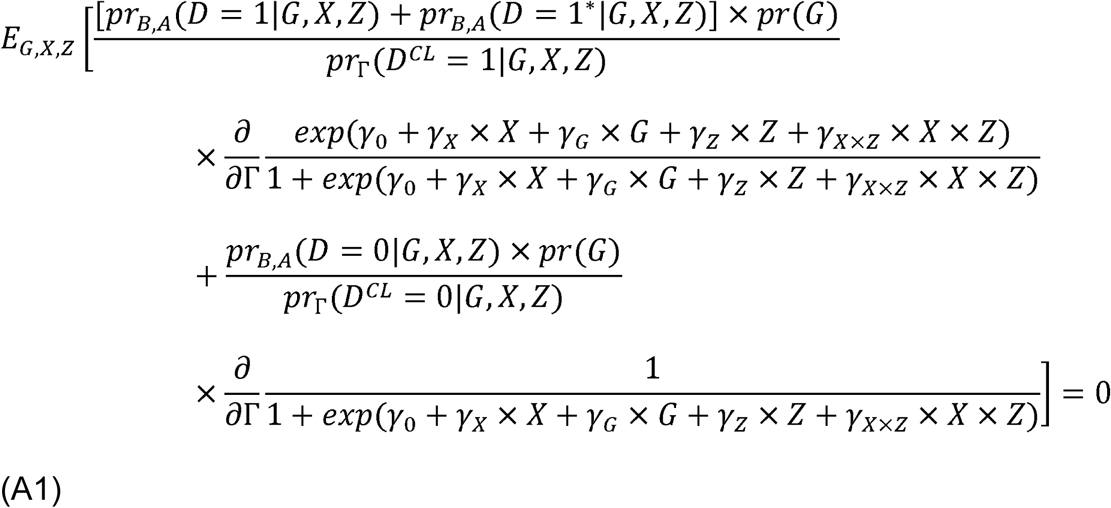

Define 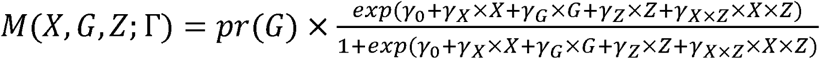

Then (A1) becomes

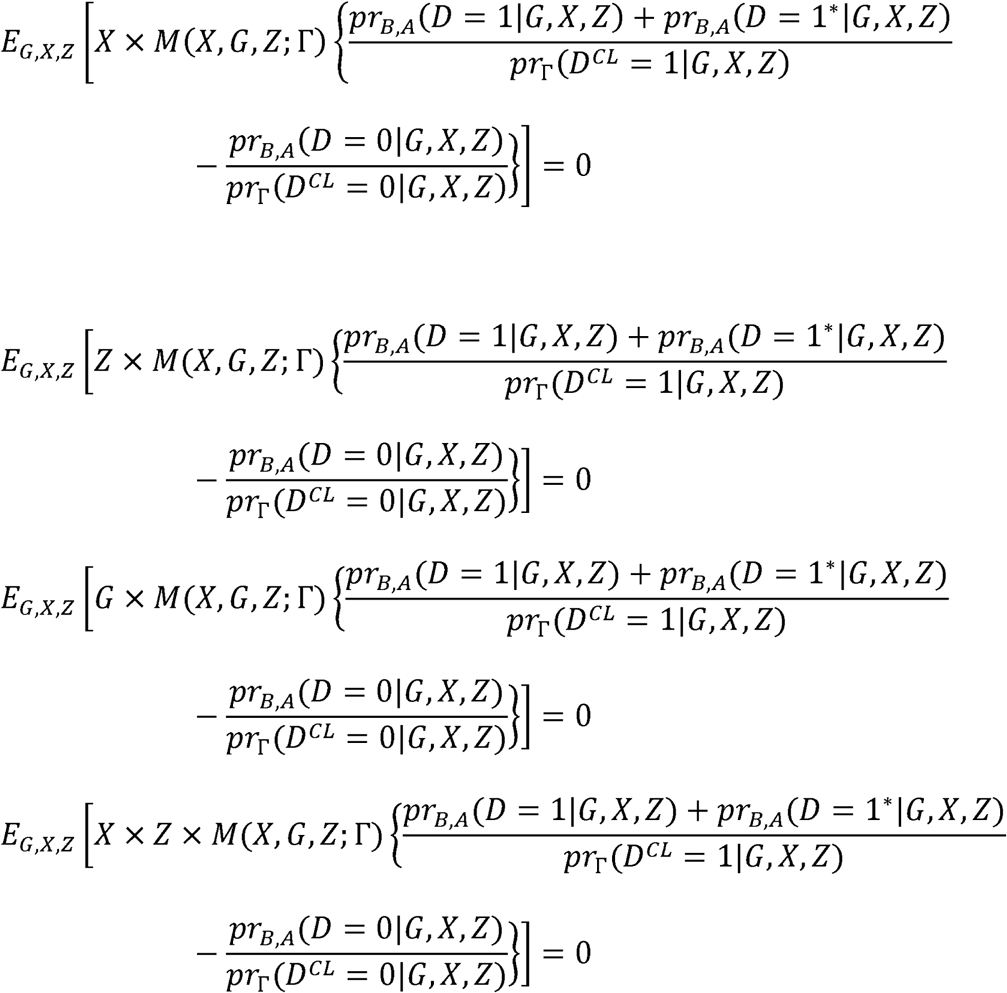

Values of Γ such that

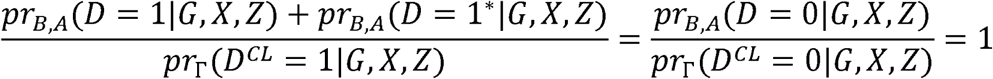

for all *G,X,Z* solve the system of equations (A1).

By definition,

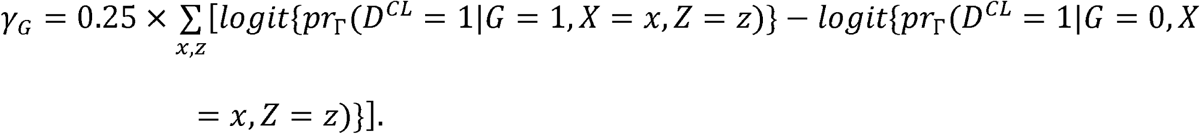

With Taylor series expansion around *β*_*G*_ =0 we arrive at (12a). Derivation for the other parameters is similar. If *X* is continuous, then e.g.,

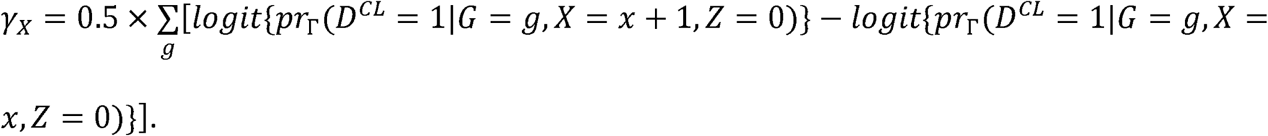

