## Supplementary material for "A simple approximation to bias in the genetic effect estimates when multiple disease states share a clinical diagnosis"

**WEB-BASED SUPPLEMENT for “A simple approximation to bias in the genetic effect estimates when multiple disease states share a clinical diagnosis”**

Iryna Lobach^1*^, Inyoung Kim^2^, Alexander Alekseyenko^3^, Siarhei Lobach^4^, Li Zhang^5^

**A2. Approximation with known** $\boldsymbol{\tau(X)}$

Suppose a researcher estimated $\gamma_{0},\gamma_{G}$ from the usual logistic regression and a reliable population estimates of $\tau(X)$ are available in the literature. Note that $\tau(X)$ can be rewritten as $\tau(X)=\frac{pr(D=1|X)}{pr(D=1|X)+pr(D=1^{*}|X)}$, hence $pr(D=1^{*}|X)=\frac{1-\tau(X)}{\tau(X)}\times pr(D=1|X)$. Then for Model 1

$\gamma_{G}\approx\log\left\{ \frac{\exp\left( \beta_{0}+\beta_{G} \right)}{\tau\left( X \right)+\left\{ \tau\left( X \right)-1 \right\}\times\exp\left( \beta_{0}+\beta_{G} \right)+\tau\left( X \right)\times\exp\left( \alpha_{0}+\alpha_{G} \right)} \right\}-\log\left\{ \frac{\exp\left( \beta_{0} \right)}{\tau\left( X \right)+\left\{ \tau\left( X \right)-1 \right\}\times\exp\left( \beta_{0} \right)+\tau\left( X \right)\times\exp\left( \alpha_{0} \right)} \right\}$ (A2)

while equation (A2) still requires $\alpha_{0}$ and $\alpha_{G}$.

For example, for Model 3, the following equations characterize the relationship between the parameters

$$\gamma_{G}\approx0.25\times\underset{x,z}{\sum}\frac{exp(\beta_{0}+\beta_{G}+\beta_{x}\times x+\beta_{Z}\times z+\beta_{X\times Z}\times x\times z)}{\tau(X)+\{\tau(X)-1\}\times exp(\beta_{0}+\beta_{G}+\beta_{x}\times x+\beta_{Z}\times z+\beta_{X\times Z}\times x\times z)+\tau(X)\times\exp(\alpha_{0}+\alpha_{G}+\alpha_{X}\times x+\alpha_{Z}\times z+\alpha_{X\times Z}\times x\times z)}$$

$$-0.25\times\underset{x,z}{\sum}\frac{exp(\beta_{0}+\beta_{x}\times x+\beta_{Z}\times z+\beta_{X\times Z}\times x\times z)}{\tau(X)+\{\tau(X)-1\}\times exp(\beta_{0}+\beta_{x}\times x+\beta_{Z}\times z+\beta_{X\times Z}\times x\times z)+\tau(X)\times\exp(\alpha_{0}+\alpha_{X}\times x+\alpha_{Z}\times z+\alpha_{X\times Z}\times x\times z)}$$

$$\gamma_{X}\approx0.5\times\underset{g}{\sum}\frac{exp(\beta_{0}+\beta_{G}\times g+\beta_{X})}{\tau(1)+\{\tau(1)-1\}\times exp(\beta_{0}+\beta_{G}\times g+\beta_{X})+\tau(1)\times\exp(\alpha_{0}+\alpha_{G}\times g+\alpha_{X})}$$

$-0.5\times\underset{g}{\sum}\frac{exp(\beta_{0}+\beta_{G}\times g)}{\tau(0)+\{\tau(0)-1\}\times exp(\beta_{0}+\beta_{G}\times g)+\tau(0)\times\exp(\alpha_{0}+\alpha_{G}\times g)}$;

$$\gamma_{Z}\approx0.5\times\underset{g}{\sum}\frac{exp(\beta_{0}+\beta_{G}\times g+\beta_{Z})}{\tau(0)+\{\tau(0)-1\}\times exp(\beta_{0}+\beta_{G}\times g+\beta_{Z})+\tau(0)\times\exp(\alpha_{0}+\alpha_{G}\times g+\alpha_{Z})}$$

$-0.5\times\underset{g}{\sum}\frac{exp(\beta_{0}+\beta_{G}\times g)}{\tau(0)+\{\tau(0)-1\}\times exp(\beta_{0}+\beta_{G}\times g)+\tau(0)\times\exp(\alpha_{0}+\alpha_{G}\times g)}$;

$$\gamma_{X\times Z}\approx0.5\times\underset{g}{\sum}\frac{exp(\beta_{0}+\beta_{G}\times g+\beta_{X}+\beta_{Z}+\beta_{X\times Z})}{\tau(0)+\{\tau(0)-1\}\times exp(\beta_{0}+\beta_{G}\times g+\beta_{X}+\beta_{Z}+\beta_{X\times Z})+\tau(0)\times\exp(\alpha_{0}+\alpha_{G}\times g+\alpha_{X}+\alpha_{Z}+\alpha_{X\times Z})}$$

$$-0.5\times\underset{g}{\sum}\frac{exp(\beta_{0}+\beta_{G}\times g+\beta_{X})}{\tau(0)+\{\tau(0)-1\}\times exp(\beta_{0}+\beta_{G}\times g+\beta_{X})+\tau(0)\times\exp(\alpha_{0}+\alpha_{G}\times g+\alpha_{X})}$$

$$-0.5\times\underset{g}{\sum}\frac{exp(\beta_{0}+\beta_{G}\times g+\beta_{Z})}{\tau(0)+\{\tau(0)-1\}\times exp(\beta_{0}+\beta_{G}\times g+\beta_{Z})+\tau(0)\times\exp(\alpha_{0}+\alpha_{G}\times g+\alpha_{Z})}$$

$+0.5\times\underset{g}{\sum}\frac{exp(\beta_{0}+\beta_{G}\times g)}{\tau(0)+\{\tau(0)-1\}\times exp(\beta_{0}+\beta_{G}\times g)+\tau(0)\times\exp(\alpha_{0}+\alpha_{G}\times g)}$ .

We note that the equations still involve the nuisance coefficients.

**A3. Model 1 when** $\boldsymbol{\tau(X)}$ **, pr(**$\boldsymbol{D}^{\boldsymbol{CL}}\boldsymbol{=1}$**) and** $\mathbf{pr(G=1)}$ **are known**

Consider Model 1. Suppose that was is reliably available are estimates of $\tau=\tau\left( X=0 \right)\times pr\left( X=0 \right)+\tau\left( X=1 \right)\times pr\left( X=1 \right)$, $pr\left( D^{CL}=1 \right)=\pi_{d^{CL}=1},$and frequency of minor allele$\theta$. Then consider the following system of equations

$$\gamma_{G}\approx log\left[ \frac{\exp\left( \beta_{0}+\beta_{G} \right)}{\tau\left( X \right)+\left\{ \tau\left( X \right)-1 \right\}\times\exp\left( \beta_{0}+\beta_{G} \right)+\exp\left( \alpha_{0}+\alpha_{G} \right)} \right]-log\left[ \frac{\exp\left( \beta_{0} \right)}{\tau\left( X \right)+\left\{ \tau\left( X \right)-1 \right\}\times\exp\left( \beta_{0} \right)+\exp\left( \alpha_{0} \right)} \right]$$

$$\gamma_{0}\approx log\left\{ \exp\left( \alpha_{0} \right)+\exp\left( \beta_{0} \right) \right\}$$

$$\pi_{d^{CL}=1}=\frac{\exp\left( \beta_{0} \right)}{1+\exp\left( \beta_{0} \right)+\exp\left( \alpha_{0} \right)}\times\left( 1-\theta\right)+\frac{\exp\left( \beta_{0}+\beta_{G} \right)}{1+\exp\left( \beta_{0}+\beta_{G} \right)+\exp\left( \alpha_{0}+\alpha_{G} \right)}\times\theta$$

$$\tau=\frac{\exp\left( \beta_{0}+\beta_{G} \right)}{1+\exp\left( \beta_{0}+\beta_{G} \right)+\exp\left( \alpha_{0}+\alpha_{G} \right)}$$

This system of equations can be solved for the parameters of interest $\beta_{0},\beta_{G}$ and the nuisance parameters $\alpha_{0},\alpha_{G}$.

Define $c=\log\left[ \frac{(\pi_{d^{CL}=1}-\tau\times\theta)\times\left\{ 1+\exp\left( \gamma_{0} \right) \right\}}{\tau\times\left( 1-\theta\right)+(\tau-2)\times(\pi_{d^{CL}=1}-\tau\times\theta)\times\left\{ 1+\exp\left( \gamma_{0} \right) \right\}+\exp\left( \gamma_{0} \right)} \right]$, then

$\alpha_{0}=\log\left[ \exp\left( \gamma_{0} \right)-\frac{(\pi_{d^{CL}=1}-\tau\times\theta)\times\left\{ 1+\exp\left( \gamma_{0} \right) \right\}}{1-\theta} \right]$,

$\beta_{0}=\log\left[ \frac{(\pi_{d^{CL}=1}-\tau\times\theta)\times\left\{ 1+\exp\left( \gamma_{0} \right) \right\}}{1-\theta} \right]$,

$\beta_{G}=\log\left\{ \frac{\tau\times c\left( \alpha_{0},\beta_{0},\tau\right)\times\exp\left( \gamma_{G} \right)}{\frac{(\tau-1)^{2}}{\tau}\times c\left( \alpha_{0},\beta_{0},\tau\right)\times\exp\left( \gamma_{G} \right)-1} \right\}-\log\left[ \frac{(\pi_{d^{CL}=1}-\tau\times\theta)\times\left\{ 1+\exp\left( \gamma_{0} \right) \right\}}{1-\theta} \right]$;

$\alpha_{G}=\log\left( \frac{1-\tau}{\tau} \right)+\beta_{0}+\beta_{G}-\alpha_{0}$.

**A4. Model 3:** $\boldsymbol{\beta}_{\boldsymbol{G}}$**,** $\boldsymbol{\beta}_{\boldsymbol{X}}$**,** $\boldsymbol{\beta}_{\boldsymbol{Z}}$**, and** $\boldsymbol{\beta}_{\boldsymbol{X\times Z}}$

We suppose that the true disease risk model is

$log\left\{ \frac{pr_{B,A}(D=1|G,X,Z)}{pr_{B,A}(D=0|G,X,Z)} \right\}=\beta_{0}+\beta_{X}\times X+\beta_{G}\times G+\beta_{Z}\times Z+\beta_{X\times Z}\times X\times Z$; (A3)

$log\left\{ \frac{pr_{B,A}(D=1^{*}|G,X,Z)}{pr_{B,A}(D=0|G,X,Z)} \right\}=\alpha_{0}+\alpha_{X}\times X+\alpha_{G}\times G+\alpha_{Z}\times Z+\alpha_{X\times Z}\times X\times Z.$	(A4)

And a model with the clinical diagnosis as an outcome variable is

$logit\left\{ pr_{\Gamma}(D^{CL}=1|G,X,Z) \right\}=\gamma_{0}+\gamma_{X}\times X+\gamma_{G}\times G+\gamma_{Z}\times Z+\gamma_{X\times Z}\times X\times Z$	(A5)

Derivations provided in Appendix A1 show that

$$\gamma_{G}\approx0.25\times\underset{x,z}{\sum}log\left( exp\{\beta_{0}+\beta_{G}+\beta_{X}\times x+\beta_{Z}\times z+\beta_{X\times Z}\times x\times z\}+exp\{\alpha_{0}+\alpha_{G}+\alpha_{X}\times x+\alpha_{Z}\times z+\alpha_{X\times Z}\times x\times z\} \right)$$

$-0.25\times\underset{x,z}{\sum}log\left( exp\{\beta_{0}+\beta_{X}\times x+\beta_{Z}\times z+\beta_{X\times Z}\times x\times z\}+exp\{\alpha_{0}+\alpha_{X}\times x+\alpha_{Z}\times z+\alpha_{X\times Z}\times x\times z\} \right)$

$$\approx0.25\times\underset{x,z}{\sum}\left\{ exp(\beta_{0}+\beta_{X}\times X+\beta_{Z}\times Z+\beta_{X\times Z}\times X\times Z)+exp(\alpha_{0}+\alpha_{G}+\alpha_{X}\times X+\alpha_{Z}\times Z+\alpha_{X\times Z}\times X\times Z) \right\}-\left\{ exp(\beta_{0}+\beta_{X}\times X+\beta_{Z}\times Z+\beta_{X\times Z}\times X\times Z)+exp(\alpha_{0}+\alpha_{X}\times X+\alpha_{Z}\times Z+\alpha_{X\times Z}\times X\times Z) \right\}$$

$+0.25\times\underset{x,z}{\sum}\frac{exp\left( \beta_{0}+\beta_{X}\times X+\beta_{Z}\times Z+\beta_{X\times Z}\times x\times z \right)}{exp\left( \beta_{0}+\beta_{X}\times X+\beta_{Z}\times Z+\beta_{X\times Z}\times x\times z \right)+exp\left( \alpha_{0}+\alpha_{G}+\alpha_{X}\times X+\alpha_{Z}\times Z+\alpha_{X\times Z}\times x\times z \right)}\times\beta_{G}$; (A6a)

$$\gamma_{X}\approx0.5\times\underset{g}{\sum}\left[ log\{exp(\beta_{0}+\beta_{G}\times g+\beta_{X})+exp(\alpha_{0}+\alpha_{G}\times g+\alpha_{X})\}-log\{exp(\beta_{0}+\beta_{G}\times g)+exp(\alpha_{0}+\alpha_{G}\times g)\} \right]$$

$\approx0.5\times\underset{g}{\sum}\left[ log\{exp(\beta_{0}+\beta_{G}\times g)+exp(\alpha_{0}+\alpha_{G}\times g+\alpha_{X})\}-log\{exp(\beta_{0}+\beta_{G}\times g)+exp(\alpha_{0}+\alpha_{X}+\alpha_{G}\times g)\}+\frac{exp(\beta_{0}+\beta_{G}\times g)}{exp(\beta_{0}+\beta_{G}\times g)+exp(\alpha_{0}+\alpha_{G}\times g+\alpha_{X})}\times\beta_{X} \right]$; (A6b)

$$\gamma_{Z}\approx0.5\times\underset{g}{\sum}\left[ log\{exp(\beta_{0}+\beta_{G}\times g+\beta_{Z})+exp(\alpha_{0}+\alpha_{G}\times g+\alpha_{Z})\}-log\{exp(\beta_{0}+\beta_{G}\times g)+exp(\alpha_{0}+\alpha_{G}\times g)\} \right]$$

$\approx0.5\times\underset{g}{\sum}\left[ log\left\{ exp\left( \beta_{0}+\beta_{G}\times g \right)+exp\left( \alpha_{0}+\alpha_{G}\times g+\alpha_{Z} \right) \right\}-log\left\{ \exp\left( \beta_{0}+\beta_{G}\times g \right)+\exp\left( \alpha_{0}+\alpha_{G}\times g \right) \right\}+\frac{\exp\left( \beta_{0}+\beta_{G}\times g \right)}{\exp\left( \beta_{0}+\beta_{G}\times g \right)+\exp\left( \alpha_{0}+\alpha_{G}\times g+\alpha_{Z} \right)}\times\beta_{Z} \right];$	(A6c)

$\gamma_{X\times Z}\approx0.5\times\underset{g}{\sum}\left[ log\left\{ exp(\beta_{0}+\beta_{G}\times g+\beta_{Z}+\beta_{X}+\beta_{X\times Z})+exp(\alpha_{0}+\alpha_{G}\times g+\alpha_{Z}+\alpha_{X}+\alpha_{X\times Z}) \right\}-log\left\{ exp(\beta_{0}+\beta_{G}\times g+\beta_{X})+exp(\alpha_{0}+\alpha_{G}\times g+\alpha_{X}) \right\}-log\left\{ exp(\beta_{0}+\beta_{G}\times g+\beta_{Z})+exp(\alpha_{0}+\alpha_{G}\times g+\alpha_{Z}) \right\}+log\left\{ exp(\beta_{0}+\beta_{G}\times g)+exp(\alpha_{0}+\alpha_{G}\times g) \right\} \right]$

$\approx0.5\times\underset{g}{\sum}\left[ log\left\{ \exp\left( \beta_{0}+\beta_{G}\times g+\beta_{Z}+\beta_{X} \right)+\exp\left( \alpha_{0}+\alpha_{G}\times g+\alpha_{Z}+\alpha_{X}+\alpha_{X\times Z} \right) \right\}-log\left\{ \exp\left( \beta_{0}+\beta_{G}\times g+\beta_{X} \right)+\exp\left( \alpha_{0}+\alpha_{G}\times g+\alpha_{X} \right) \right\}-log\left\{ \exp\left( \beta_{0}+\beta_{G}\times g+\beta_{Z} \right)+\exp\left( \alpha_{0}+\alpha_{G}\times g+\alpha_{Z} \right) \right\}+log\left\{ \exp\left( \beta_{0}+\beta_{G}\times g \right)+\exp\left( \alpha_{0}+\alpha_{G}\times g \right) \right\}+\frac{\exp\left( \beta_{0}+\beta_{G}\times g+\beta_{Z}+\beta_{X} \right)}{\exp\left( \beta_{0}+\beta_{G}\times g+\beta_{Z}+\beta_{X} \right)+\exp\left( \alpha_{0}+\alpha_{G}\times g+\alpha_{Z}+\alpha_{X}+\alpha_{X\times Z} \right)}\times\beta_{X\times Z} \right];$	(A6d)

Remarks:

1. Model 3, equation (A6a). If $\beta_{G}=\alpha_{G}=0$, then $\gamma_{G}=0.$

2. Model 3, equation (A6b). If $\beta_{X}=\alpha_{X}=0$, then $\gamma_{X}=0.$

3. Model 3, equation (A6v). If $\beta_{Z}=\alpha_{Z}=0$, then $\gamma_{Z}=0.$

4. Model 3, equation (A6d). If $\beta_{Z}=\beta_{X}=\beta_{X\times Z}=0$, then $\gamma_{X\times Z}=0.$

**A5.** **Model 4:** $\boldsymbol{\beta}_{\boldsymbol{G}_{\boldsymbol{1}}}\boldsymbol{,}\boldsymbol{\beta}_{\boldsymbol{G}_{\boldsymbol{2}}}$ **and** $\boldsymbol{\beta}_{\boldsymbol{G}_{\boldsymbol{1}}\boldsymbol{\times}\boldsymbol{G}_{\boldsymbol{2}}}$**.** We next consider a setting when the genetic variable $G$ are in the risk model, i.e. the true disease risk model is

$\log\left\{ \frac{pr_{B,A}(D=1|G_{1},G_{2})}{pr_{B,A}(D=0|G_{1},G_{2})} \right\}=\beta_{0}+\beta_{G_{1}}\times G_{1}+\beta_{G_{2}}\times G_{2}+\beta_{G_{1}\times G_{2}}\times G_{1}\times G_{2},$	(A7)

$\log\left\{ \frac{pr_{B,A}(D=1^{*}|G_{1},G_{2})}{pr_{B,A}(D=0|G_{1},G_{2})} \right\}=\alpha_{0}+\alpha_{G_{1}}\times G_{1}+\alpha_{G_{2}}\times G_{2}+\alpha_{G_{1}\times G_{2}}\times G_{1}\times G_{2};$	(A8)

while the model used is

$\log\left\{ \frac{pr_{\Gamma}(D^{CL}=1|G_{1},G_{2})}{pr_{\Gamma}(D^{CL}=0|G_{1},G_{2})} \right\}=\gamma_{0}+\gamma_{G_{1}}\times G_{1}+\gamma_{G_{2}}\times G_{2}+\gamma_{G_{1}\times G_{2}}\times G_{1}\times G_{2}.$	(A9)

Derivations provided in Appendix A2 show that

$\gamma_{0}\approx\log\left\{ \exp\left( \beta_{0} \right)+\exp\left( \alpha_{0} \right) \right\};$	(A10a)

$\gamma_{G_{1}}\approx0.5\times\sum_{g_{2}} \left[ \log\left\{ \exp\left( \beta_{0}+\beta_{G_{1}}+\beta_{G_{2}}\times g_{2}+\beta_{G_{1}\times G_{2}}\times g_{2} \right)+\exp\left( \alpha_{0}+\alpha_{G_{1}}+\alpha_{G_{2}}\times g_{2}+\alpha_{g_{1}\times g_{2}}\times g_{2} \right) \right\}-log\{exp(\beta_{0}+\beta_{G_{2}}\times g_{2}+\beta_{G_{1}\times G_{2}}\times g_{2})+exp(\alpha_{0}+\alpha_{G_{2}}\times g_{2}+\alpha_{g_{1}\times g_{2}}\times g_{2})\} \right]$

$\approx0.5\times\sum_{g_{2}} \left[ \log\left\{ \exp\left( \beta_{0}+\beta_{G_{2}}\times g_{2}+\beta_{G_{1}\times G_{2}}\times g_{2} \right)+\exp\left( \alpha_{0}+\alpha_{G_{1}}+\alpha_{G_{2}}\times g_{2}+\alpha_{g_{1}\times g_{2}}\times g_{2} \right) \right\}-\log\left\{ \exp\left( \beta_{0}+\beta_{G_{2}}\times g_{2}+\beta_{G_{1}\times G_{2}}\times g_{2} \right)+\exp\left( \alpha_{0}+\alpha_{G_{1}}+\alpha_{G_{2}}\times g_{2}+\alpha_{g_{1}\times g_{2}}\times g_{2} \right) \right\} \right] +0.5\times\sum_{g_{2}} \frac{\exp\left( \beta_{0}+\beta_{G_{2}}\times g_{2}+\beta_{G_{1}\times G_{2}}\times g_{2} \right)}{\exp\left( \beta_{0}+\beta_{G_{2}}\times g_{2}+\beta_{G_{1}\times G_{2}}\times g_{2} \right)+\exp\left( \alpha_{0}+\alpha_{G_{1}}+\alpha_{G_{2}}\times g_{2}+\alpha_{G_{1}\times G_{2}}\times g_{2} \right)}\times\beta_{G_{1}};$	(A10b)

$\gamma_{G_{2}}\approx0.5\times\sum_{g_{1}} \left[ \log\left\{ \exp\left( \beta_{0}+\beta_{G_{1}}\times g_{1}+\beta_{G_{2}}+\beta_{G_{1}\times G_{2}}\times g_{1} \right)+\exp\left( \alpha_{0}+\alpha_{G_{1}}\times g_{1}+\alpha_{G_{2}}+\alpha_{G_{1}\times G_{2}}\times g_{1} \right) \right\}-log\{exp(\beta_{0}+\beta_{G_{1}}\times g_{1}+\beta_{G_{1}\times G_{2}}\times g_{1})+exp(\alpha_{0}+\alpha_{G_{1}}\times g_{1}+\alpha_{G_{1}\times G_{2}}\times g_{1})\} \right]$

$\approx0.5\times\sum_{g_{1}} \left[ \log\left\{ \exp\left( \beta_{0}+\beta_{G_{1}}\times g_{1}+\beta_{G_{1}\times G_{2}}\times g_{1} \right)+\exp\left( \alpha_{0}+\alpha_{G_{1}}\times g_{1}+\alpha_{G_{2}}+\alpha_{g_{1}\times g_{2}}\times g_{1} \right) \right\}-\log\left\{ \exp\left( \beta_{0}+\beta_{G_{1}}\times g_{1}+\beta_{G_{1}\times G_{2}}\times g_{1} \right)+\exp\left( \alpha_{0}+\alpha_{G_{1}}\times g_{1}+\alpha_{G_{1}\times G_{2}}\times g_{2} \right) \right\} \right] +0.5\times\sum_{g_{2}} \frac{\exp\left( \beta_{0}+\beta_{G_{1}}\times g_{1}+\beta_{G_{1}\times G_{2}}\times g_{1} \right)}{\exp\left( \beta_{0}+\beta_{G_{1}}\times g_{1}+\beta_{G_{1}\times G_{2}}\times g_{1} \right)+\exp\left( \alpha_{0}+\alpha_{G_{1}}\times g_{1}+\alpha_{G_{2}}+\alpha_{G_{1}\times G_{2}}\times g_{1} \right)}\times\beta_{G_{2}};$	(A10c)

$\gamma_{{G_{1}\times G}_{2}}\approx\log\left\{ \exp\left( \beta_{0}+\beta_{G_{1}}+\beta_{G_{2}}+\beta_{G_{1}\times G_{2}} \right)+\exp\left( \alpha_{0}+\alpha_{G_{1}}+\alpha_{G_{2}}+\alpha_{G_{1}\times G_{2}} \right) \right\}-\log\left\{ \exp\left( \beta_{0}+\beta_{G_{1}} \right)+\exp\left( \alpha_{0}+\alpha_{G_{1}} \right) \right\}-\log\left\{ \exp\left( \beta_{0}+\beta_{G_{2}} \right)+\exp\left( \alpha_{0}+\alpha_{G_{2}} \right) \right\}+\log\left\{ \exp\left( \beta_{0} \right)+\exp\left( \alpha_{0} \right) \right\}$	(A10d)

**Figure 1:** Empirical biases in estimates of $\beta_{G},\beta_{A},\beta_{S},\beta_{\epsilon4},\beta_{A\times\epsilon4}$ across 500 datasets with 3,000 cases and 3,000 controls when the disease states are simulated according to risk model (1)-(2), but the parameters are estimated using model (3). Here the genotype, ApoE $\epsilon4$ status and $A\times\epsilon4$ are not associated with the nuisance disease state. Genotype, age, sex, ApoE $\epsilon4$ status are simulated to be binary with frequencies $\theta_{G}=0.10,\theta_{A}=0.50,\theta_{S}=0.52,\theta_{\epsilon4}=0.07$**.** The disease states are then simulated according to risk model (1)-(2) with $\beta_{0}=-1,\beta_{S}=log(0.92)=-0.08,\beta_{\epsilon4}=log(8)=2.1,\beta_{A}=log(2)=0.69,$ $\beta_{G}=log(1),log(1.5),log(2),log(2.5),log(3),\beta_{A\times\epsilon4}=log\left( 1 \right)=0,log\left( 1.5 \right)=0.41,log\left( 2 \right)=0.69,log\left( 2.5 \right)=0.92,$ $\alpha_{0}=-1,\alpha_{S}=log\left( 0.92 \right),\alpha_{\epsilon4}=0,\alpha_{A}=log\left( 2 \right),\alpha_{G}=0,\alpha_{A\times\epsilon4}=0.$ Values of $\beta_{A\times\epsilon4}$ are along the x-axis and values of $\beta_{G}$ are indicated by color.

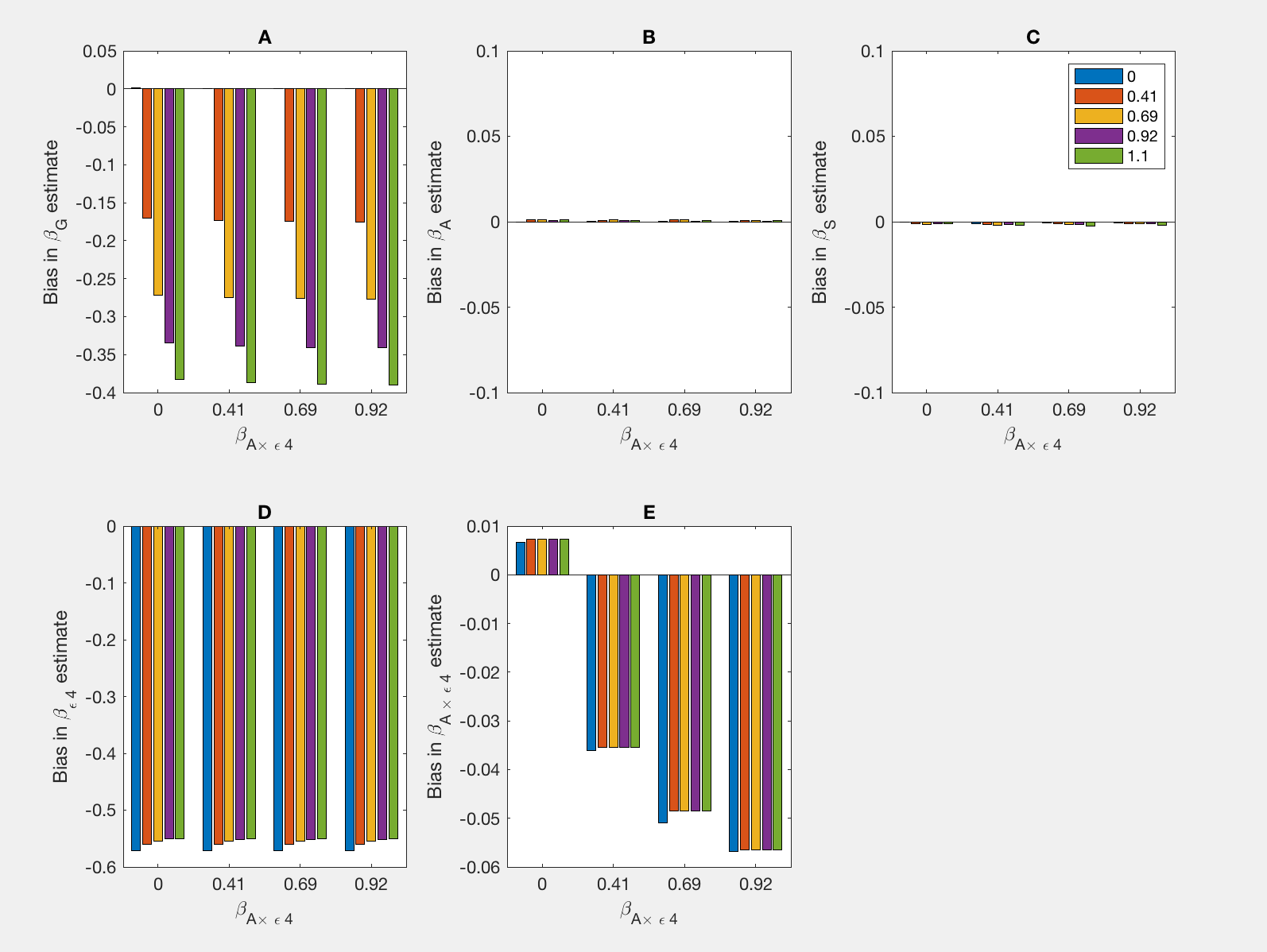

**Figure 2:** Empirical bias (Emp) and bias approximated by (A6b)(AX) in $\beta_{G}$ estimates when the data are generated according to disease states as in (A3)-(A4), but the parameters are estimated in the model (A5). Empirical estimates are the averages 500 datasets with 3,000 cases and 3,000 controls. Genotype, age, sex, ApoE $\epsilon4$ status are simulated to be binary with frequencies $\theta_{G}=0.10,\theta_{A}=0.50,\theta_{S}=0.52,\theta_{\epsilon4}=0.07$**.** The pathologically defined disease states are then simulated according to risk model (A3)-(A4) with $\beta_{0}=-1,\beta_{S}=log\left( 0.92 \right)=-0.08,\beta_{\epsilon4}=log\left( 8 \right)=2.1,\beta_{A}=log\left( 2 \right)=0.69,\beta_{G}=log\left( 1 \right)=0,log\left( 1.5 \right)=0.41,log\left( 2 \right)=0.69,log\left( 2.5 \right)=0.92,log\left( 3 \right)=1.1,\beta_{A\times\epsilon4}=log\left( 1 \right)=0,log\left( 1.5 \right)=0.41,log\left( 2 \right)=0.69,log\left( 2.5 \right)=0.92, \alpha_{0}=-1,\alpha_{S}=log\left( 0.92 \right),\alpha_{\epsilon4}=0,\alpha_{A}=log\left( 2 \right),\alpha_{G}=0,\alpha_{A\times\epsilon4}=0.$ Values of $\beta_{A\times\epsilon4}$ are along the x-axis and empirical estimates are shown in blue, approximations (A6a-c) are in red.

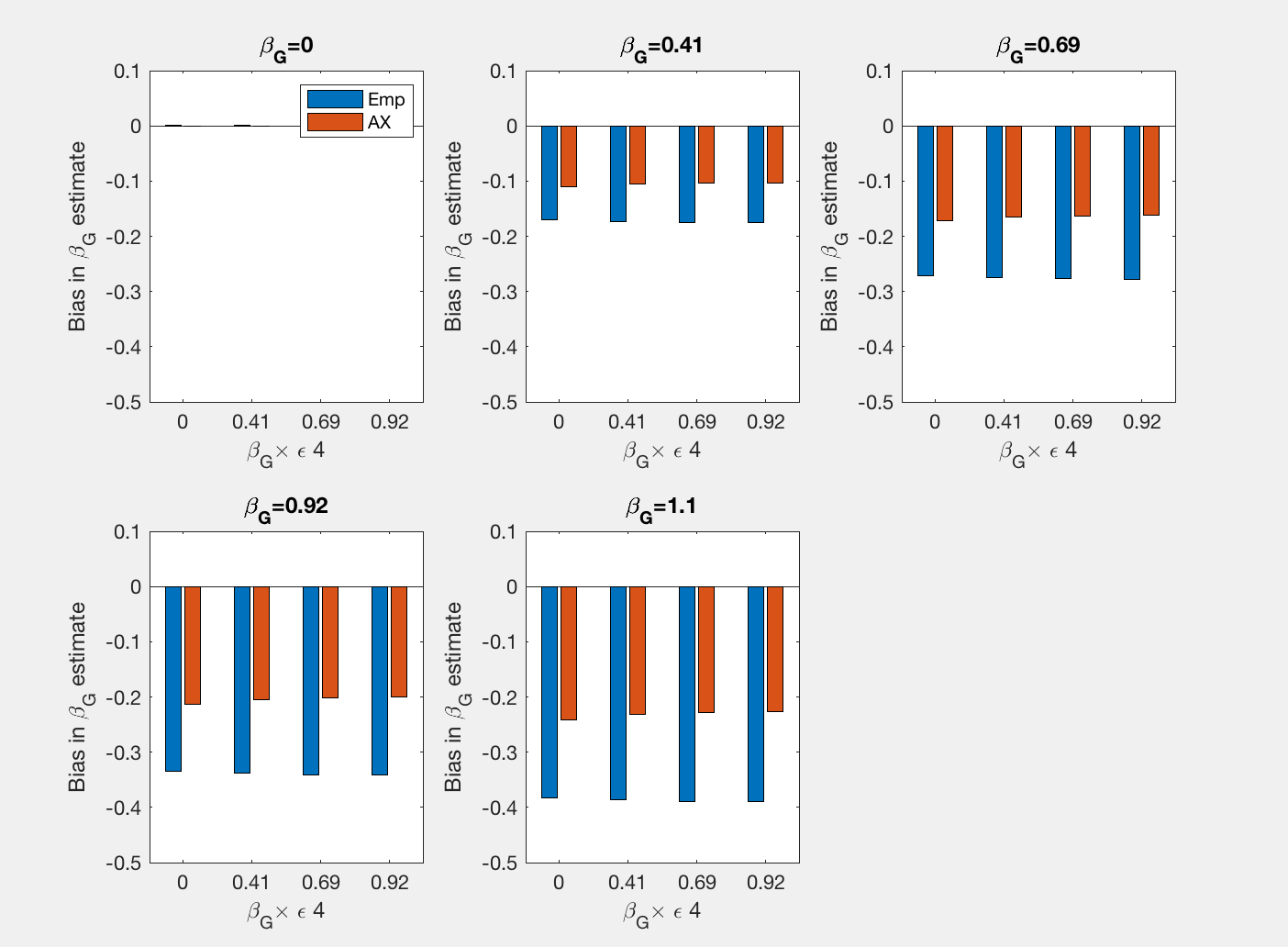

**Figure 3:** Empirical bias (Emp) and bias approximated by (A6b)(AX) in $\beta_{\epsilon4}$ estimates when the data are generated according to disease states as in (A3)-(A4), but the parameters are estimated in the model (A5). Genotype, age, sex, ApoE $\epsilon4$ status are simulated to be binary with frequencies $\theta_{G}=0.10,\theta_{A}=0.50,\theta_{S}=0.52,\theta_{\epsilon4}=0.07$**.** The pathologically defined disease states are then simulated according to risk model (A3)-(A4) $\beta_{0}=-1,\beta_{S}=log(0.92)=-0.08,\beta_{\epsilon4}=log(8)=2.1,\beta_{A}=log(2)=0.69,$ $\beta_{G}=log\left( 1 \right)=0,log\left( 1.5 \right)=0.41,log\left( 2 \right)=0.69,log\left( 2.5 \right)=0.92,log\left( 3 \right)=1.1,\beta_{A\times\epsilon4}=log\left( 1 \right)=0,log\left( 1.5 \right)=0.41,log\left( 2 \right)=0.69,log\left( 2.5 \right)=0.92,$ $\alpha_{0}=-1,\alpha_{S}=log\left( 0.92 \right),\alpha_{\epsilon4}=0,\alpha_{A}=log\left( 2 \right),\alpha_{G}=0,\alpha_{A\times\epsilon4}=0.$ Values of $\beta_{A\times\epsilon4}$ are along the x-axis and empirical estimates are shown in blue, approximations are in red..

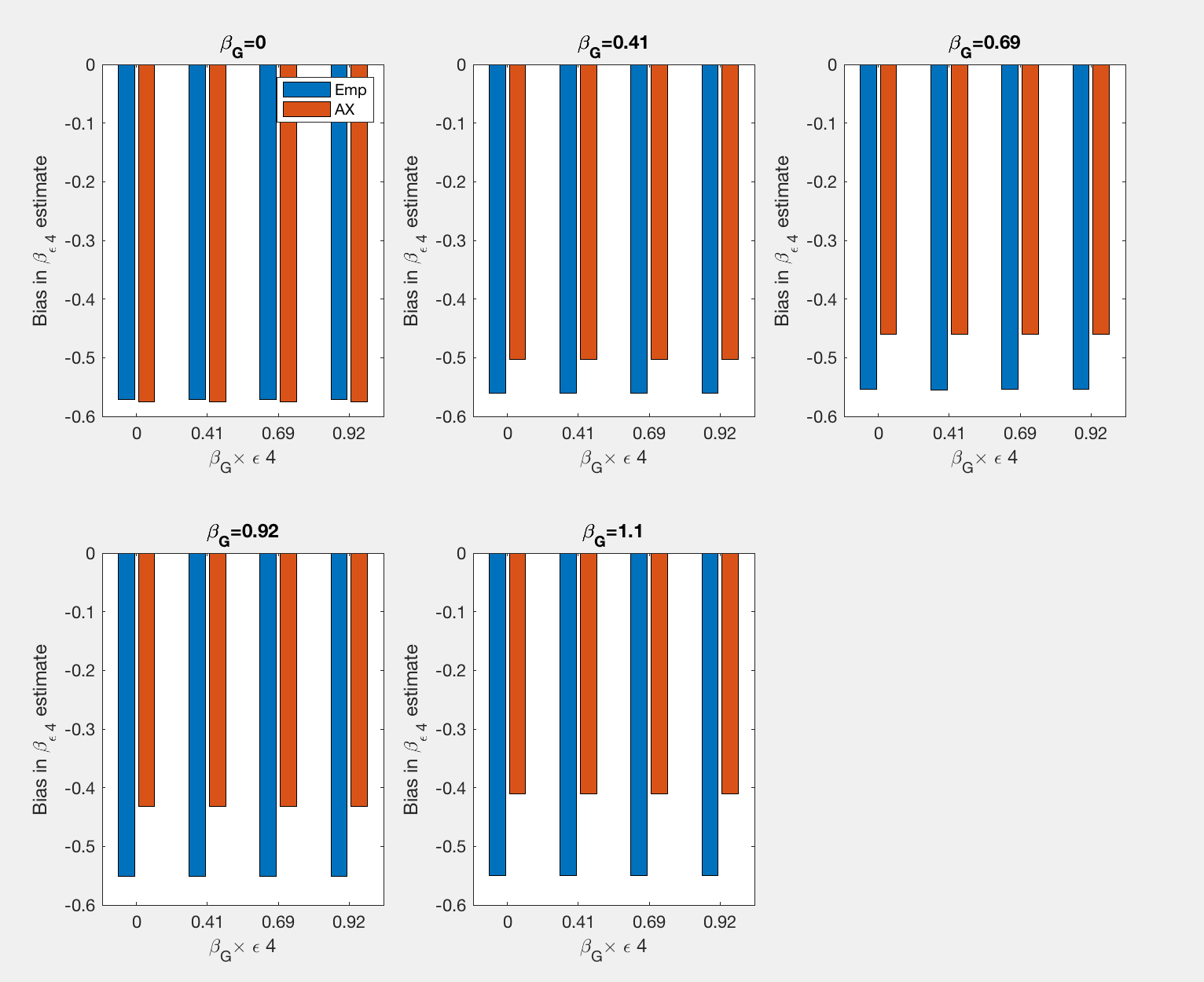

**Figure 4:** Empirical bias (Emp) and bias approximated by (A6b)(AX) in $\beta_{A\times\epsilon4}$estimates when the data are generated according to disease states as in (A3)-(A4), but the parameters are estimated in the model (A5). Genotype, age, sex, ApoE $\epsilon4$ status are simulated to be binary with frequencies $\theta_{G}=0.10,\theta_{A}=0.50,\theta_{S}=0.52,\theta_{\epsilon4}=0.07$**.** The pathologically defined disease states are then simulated according to risk model (A3)-(A4) $\beta_{0}=-1,\beta_{S}=log(0.92)=-0.08,\beta_{\epsilon4}=log(8)=2.1,\beta_{A}=log(2)=0.69,$ $\beta_{G}=log\left( 1 \right)=0,log\left( 1.5 \right)=0.41,log\left( 2 \right)=0.69,log\left( 2.5 \right)=0.92,log\left( 3 \right)=1.1,\beta_{A\times\epsilon4}=log\left( 1 \right)=0,log\left( 1.5 \right)=0.41,log\left( 2 \right)=0.69,log\left( 2.5 \right)=0.92,$ $\alpha_{0}=-1,\alpha_{S}=log\left( 0.92 \right),\alpha_{\epsilon4}=0,\alpha_{A}=log\left( 2 \right),\alpha_{G}=0,\alpha_{A\times\epsilon4}=0.$ Values of $\beta_{A\times\epsilon4}$ are along the x-axis and empirical estimates are shown in blue, approximations are in red..

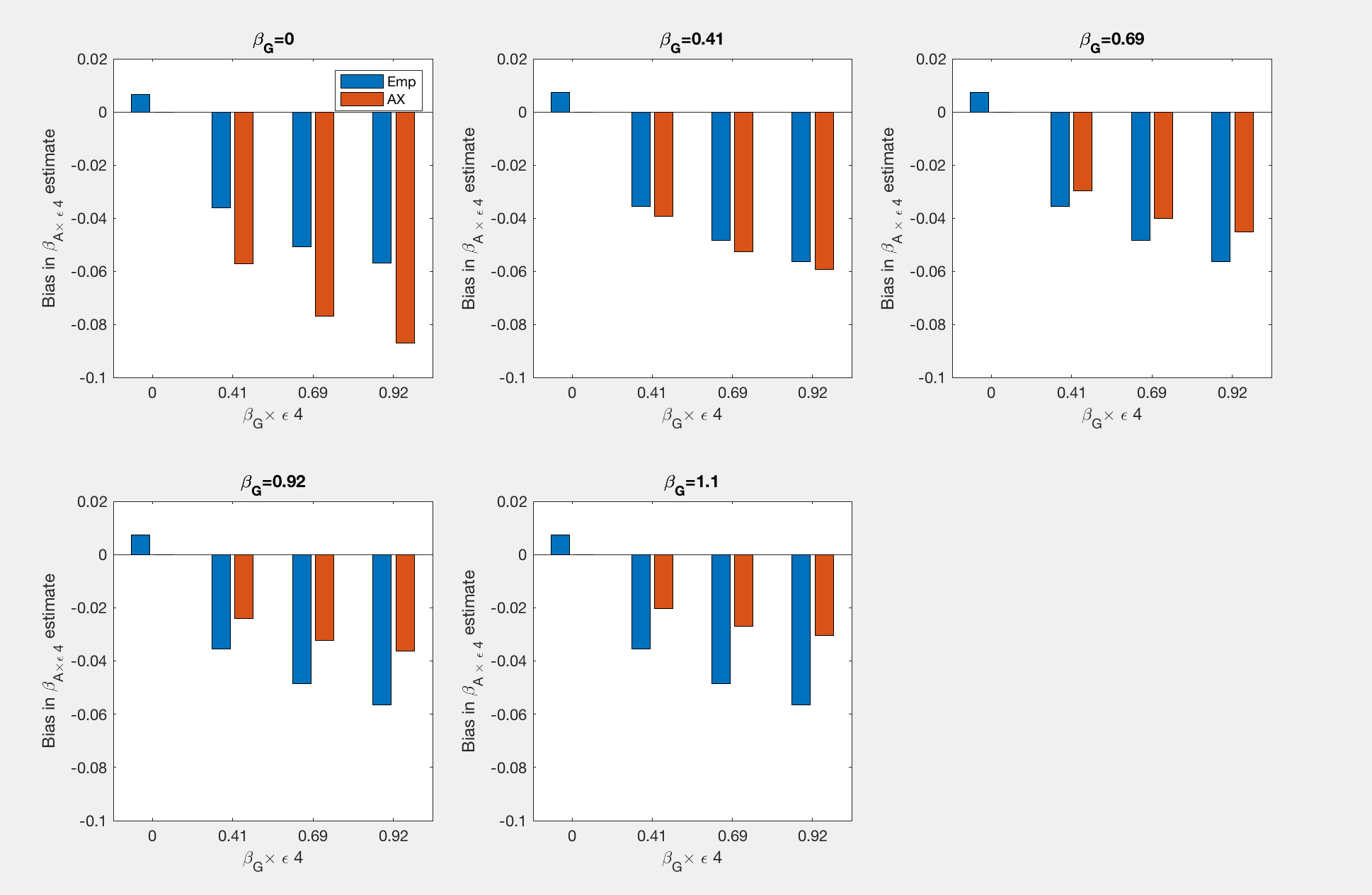

**Figure 5:** Empirical biases in estimates of $\beta_{G},\beta_{A},\beta_{S},\beta_{\epsilon4},\beta_{A\times\epsilon4}$ across 500 datasets with 3,000 cases and 3,000 controls when the disease states are generated as in (A3)-(A4), but the parameters are estimated in the model (A5). Genotype, age, sex, ApoE $\epsilon4$ status are simulated to be binary with frequencies $\theta_{G}=0.10,\theta_{A}=0.50,\theta_{S}=0.52,\theta_{\epsilon4}=0.07$**.** The pathologically defined disease states are then simulated according to risk model (1)-(2) with $\beta_{0}=1.5,\beta_{S}=log(0.80),\beta_{\epsilon4}=log(8),\beta_{A}=log(3),$

$\beta_{G}=log\left( 1 \right)=0,log\left( 1.5 \right)=0.41,log\left( 2 \right)=0.69,log\left( 2.5 \right)=0.92,log\left( 3 \right)=1.1, \beta_{A\times\epsilon4}=log\left( 1 \right)=0,log\left( 1.5 \right)=0.41,log\left( 2 \right)=0.69,log\left( 2.5 \right)=0.92, \alpha_{0}=0.5,\alpha_{S}=log(0.80),\alpha_{\epsilon4}=log(4),\alpha_{A}=log(3),\alpha_{G}=log(2),\alpha_{A\times\epsilon4}=log(2).$ Values of $\beta_{A\times\epsilon4}$ are along the x-axis and values of $\beta_{G}$ are indicated by color.

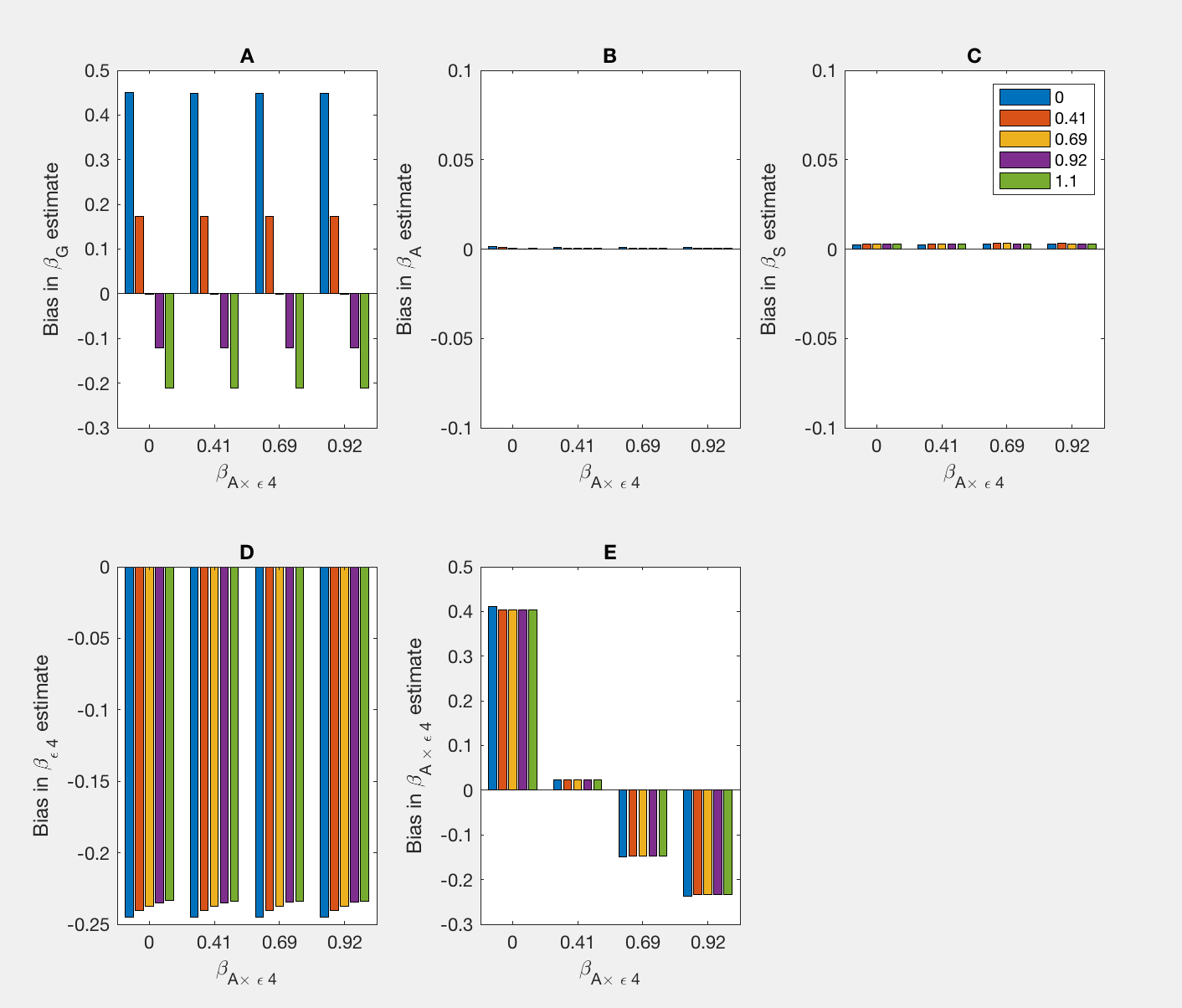

**Supplementary Figure 1:** Empirical biases in estimates of $\beta_{G},\beta_{A},\beta_{S},\beta_{\epsilon4},\beta_{A\times\epsilon4}$ across 500 datasets with 30,000 cases and 30,000 controls. Genotype, age, sex, ApoE $\epsilon4$ status are simulated to be binary with frequencies $\theta_{G}=0.10,\theta_{A}=0.50,\theta_{S}=0.52,\theta_{\epsilon4}=0.07$**.** The pathologically defined disease states are then simulated according to risk model (1)-(2) with the clinical diagnosis and disease states with coefficients $\beta_{0}=-1,\beta_{S}=log(0.92)=-0.08,\beta_{\epsilon4}=log(8)=2.1,\beta_{A}=log(2)=0.69,$

$\beta_{G}=log\left( 1 \right)=0,log\left( 1.5 \right)=0.41,log\left( 2 \right)=0.69,log\left( 2.5 \right)=0.92,log\left( 3 \right)=1.1,\beta_{A\times\epsilon4}=log\left( 1 \right)=0,log\left( 1.5 \right)=0.41,log\left( 2 \right)=0.69,log\left( 2.5 \right)=0.92,$ $\alpha_{0}=-1,\alpha_{S}=log(0.92),\alpha_{\epsilon4}=0,\alpha_{A}=log(2),\alpha_{G}=0,\alpha_{A\times\epsilon4}=0.$ Values of $\beta_{A\times\epsilon4}$ are along the x-axis and values of $\beta_{G}$ are indicated by color.
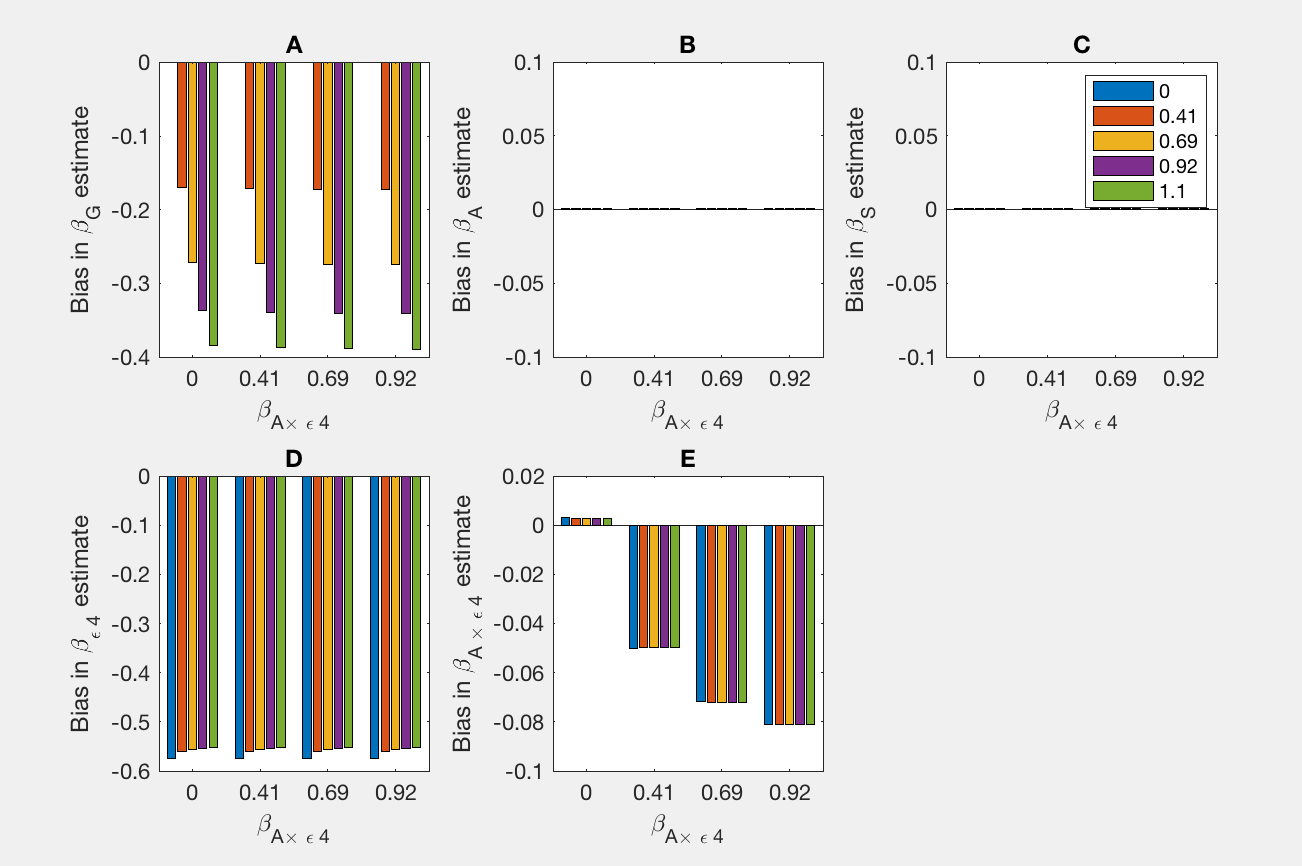

**Supplementary Figure 2:** Empirical biases in estimates of $\beta_{G},\beta_{A},\beta_{S},\beta_{\epsilon4},\beta_{A\times\epsilon4}$ across 500 datasets with 3,000 cases and 3,000 controls. Genotype, age, sex, ApoE $\epsilon4$ status are simulated to be binary with frequencies $\theta_{G}=0.10,\theta_{A}=0.50,\theta_{S}=0.52,\theta_{\epsilon4}=0.07$**.** The pathologically defined disease states are then simulated according to risk model (1)-(2) with the clinical diagnosis and disease states with coefficients $\beta_{0}=-1,\beta_{S}=log(0.92)=-0.08,\beta_{\epsilon4}=log(8)=2.1,\beta_{A}=log(2)=0.69,$

$\beta_{G}=log\left( 1 \right)=0,log\left( 1.5 \right)=0.41,log\left( 2 \right)=0.69,log\left( 2.5 \right)=0.92,log\left( 3 \right)=1.1,\beta_{A\times\epsilon4}=log\left( 1 \right)=0,log\left( 1.5 \right)=0.41,log\left( 2 \right)=0.69,log\left( 2.5 \right)=0.92,\alpha_{0}=-1,\alpha_{S}=log\left( 0.92 \right),\alpha_{\epsilon4}=log\left( 2 \right),\alpha_{A}=log\left( 2 \right),\alpha_{G}=log\left( 1.5 \right),\alpha_{A\times\epsilon4}=0.$ Values of $\beta_{A\times\epsilon4}$ are along the x-axis and values of $\beta_{G}$ are indicated by color.
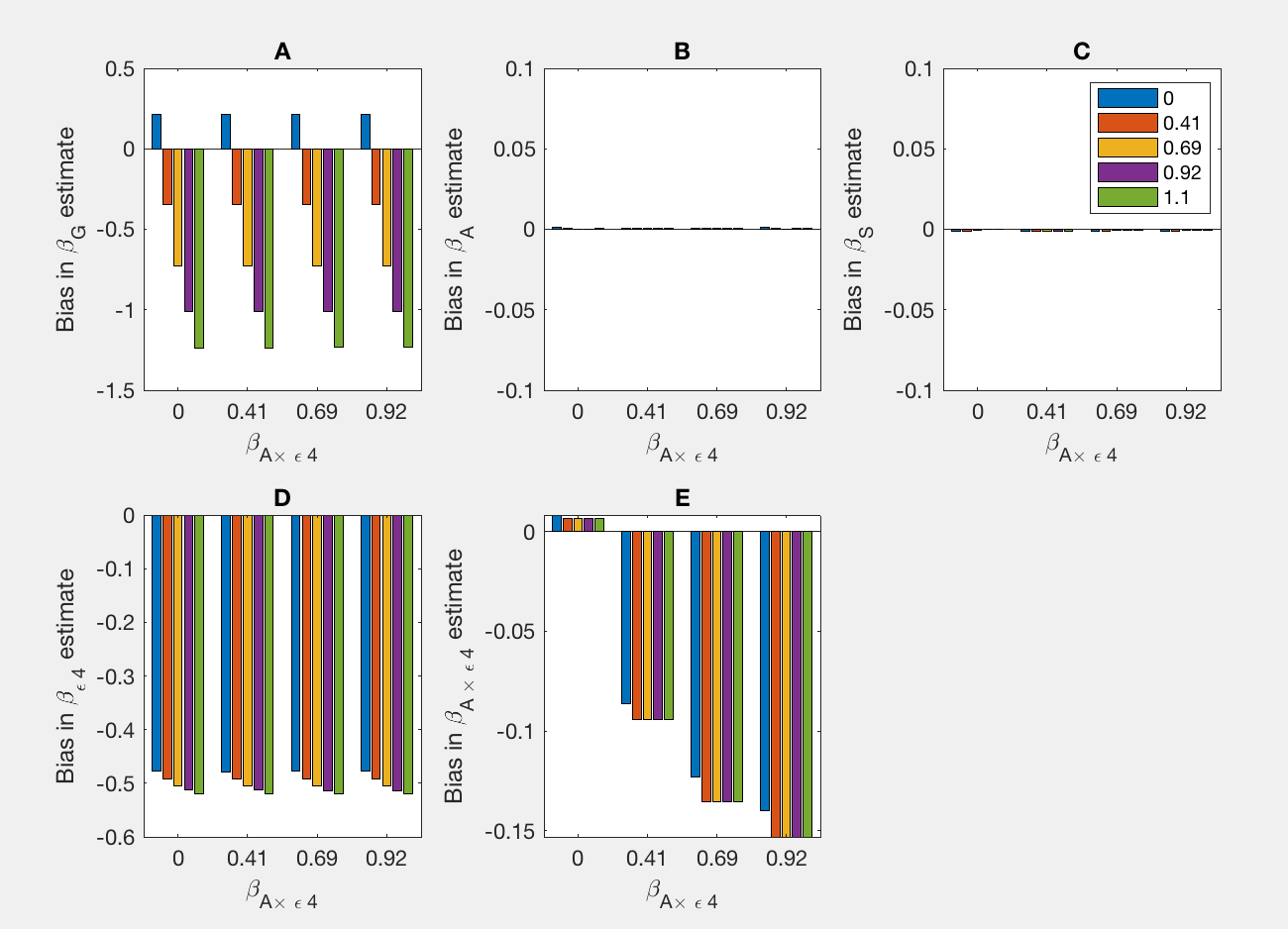

**Supplementary Figure 3:** Empirical biases in estimates of $\beta_{G},\beta_{A},\beta_{S},\beta_{\epsilon4},\beta_{A\times\epsilon4}$ across 500 datasets with 3,000 cases and 3,000 controls. Genotype, age, sex, ApoE $\epsilon4$ status are simulated to be binary with frequencies $\theta_{G}=0.10,\theta_{A}=0.50,\theta_{S}=0.52,\theta_{\epsilon4}=0.07$**.** The pathologically defined disease states are then simulated according to risk model (1)-(2) with the clinical diagnosis and disease states with coefficients $\beta_{0}=1.5,\beta_{S}=log(0.80),\beta_{\epsilon4}=log(8),\beta_{A}=log(3),$

$\beta_{G}=log\left( 1 \right)=0,log\left( 1.5 \right)=0.41,log\left( 2 \right)=0.69,log\left( 2.5 \right)=0.92,log\left( 3 \right)=1.1,\beta_{A\times\epsilon4}=log\left( 1 \right)=0,log\left( 1.5 \right)=0.41,log\left( 2 \right)=0.69,log\left( 2.5 \right)=0.92,\alpha_{0}=-1,\alpha_{S}=log(0.92),\alpha_{\epsilon4}=0,\alpha_{A}=log(2),\alpha_{G}=0,\alpha_{A\times\epsilon4}=0.$Values of $\beta_{A\times\epsilon4}$ are along the x-axis and values of $\beta_{G}$ are indicated by color.
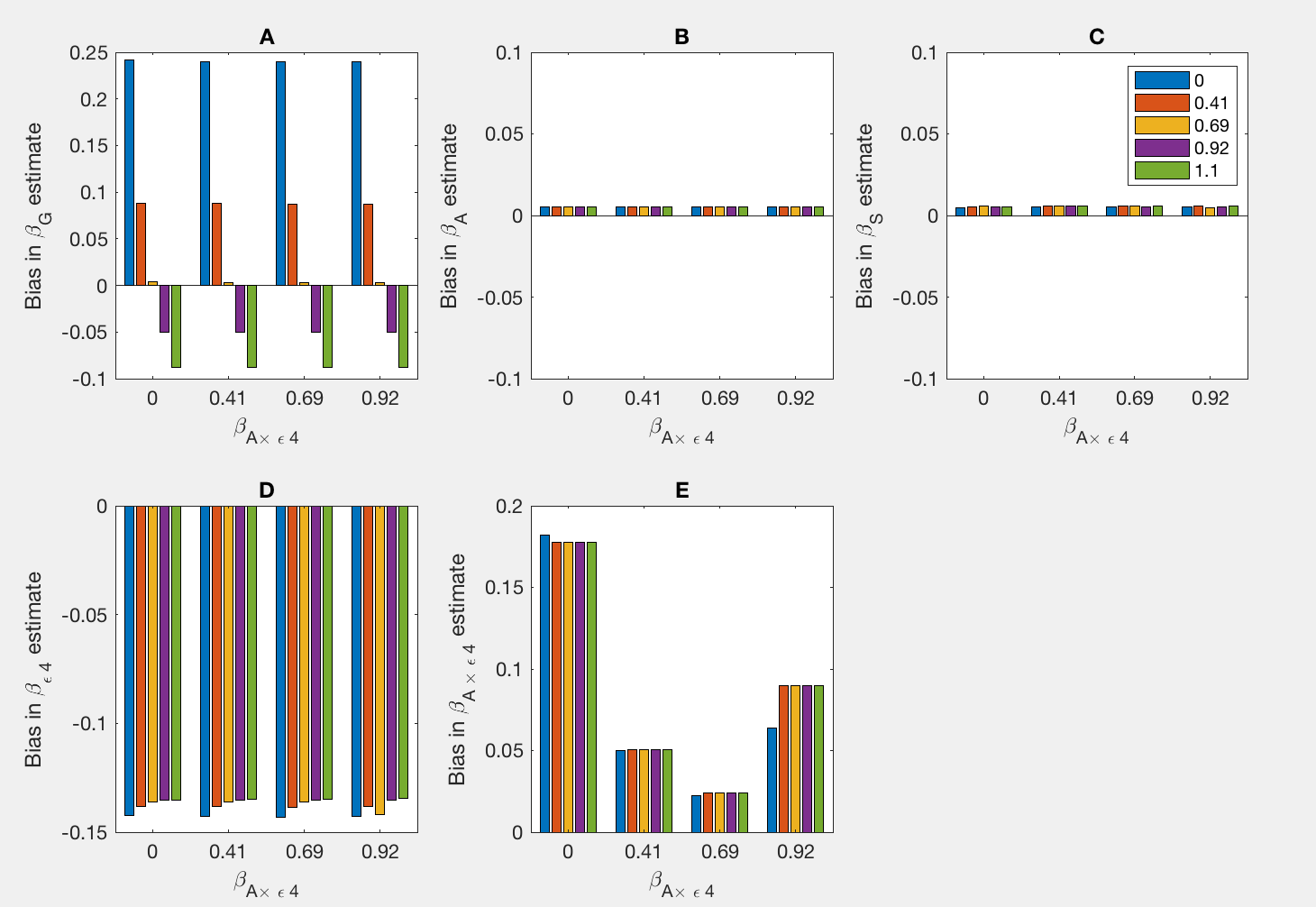

**Supplementary Figure 4:** Empirical bias (Emp) and approximation (A6b) (AX) in estimates of $\beta_{G}$ across 500 datasets with 3,000 cases and 3,000 controlswhen the data are simulated using model (A3)-(A4) but the parameters are estimated using model (A5). Genotype, age, sex, ApoE $\epsilon4$ status are simulated to be binary with frequencies $\theta_{G}=0.10,\theta_{A}=0.50,\theta_{S}=0.52,\theta_{\epsilon4}=0.07$**.** The coefficients are $\beta_{0}=1.5,\beta_{S}=log(0.80),\beta_{\epsilon4}=log(8),\beta_{A}=log(3),$

$\beta_{G}=log\left( 1 \right)=0,log\left( 1.5 \right)=0.41,log\left( 2 \right)=0.69,log\left( 2.5 \right)=0.92,log\left( 3 \right)=1.1,\beta_{A\times\epsilon4}=log\left( 1 \right)=0,log\left( 1.5 \right)=0.41,log\left( 2 \right)=0.69,log\left( 2.5 \right)=0.92,\alpha_{0}=0.5,\alpha_{S}=log\left( 0.80 \right),\alpha_{\epsilon4}=log\left( 4 \right),\alpha_{A}=log\left( 3 \right),\alpha_{G}=log\left( 2 \right),\alpha_{A\times\epsilon4}=log\left( 2 \right).$ Values of $\beta_{A\times\epsilon4}$ are along the x-axis and values of $\beta_{G}$. Empirical estimates are in blue and the approximations are in red.

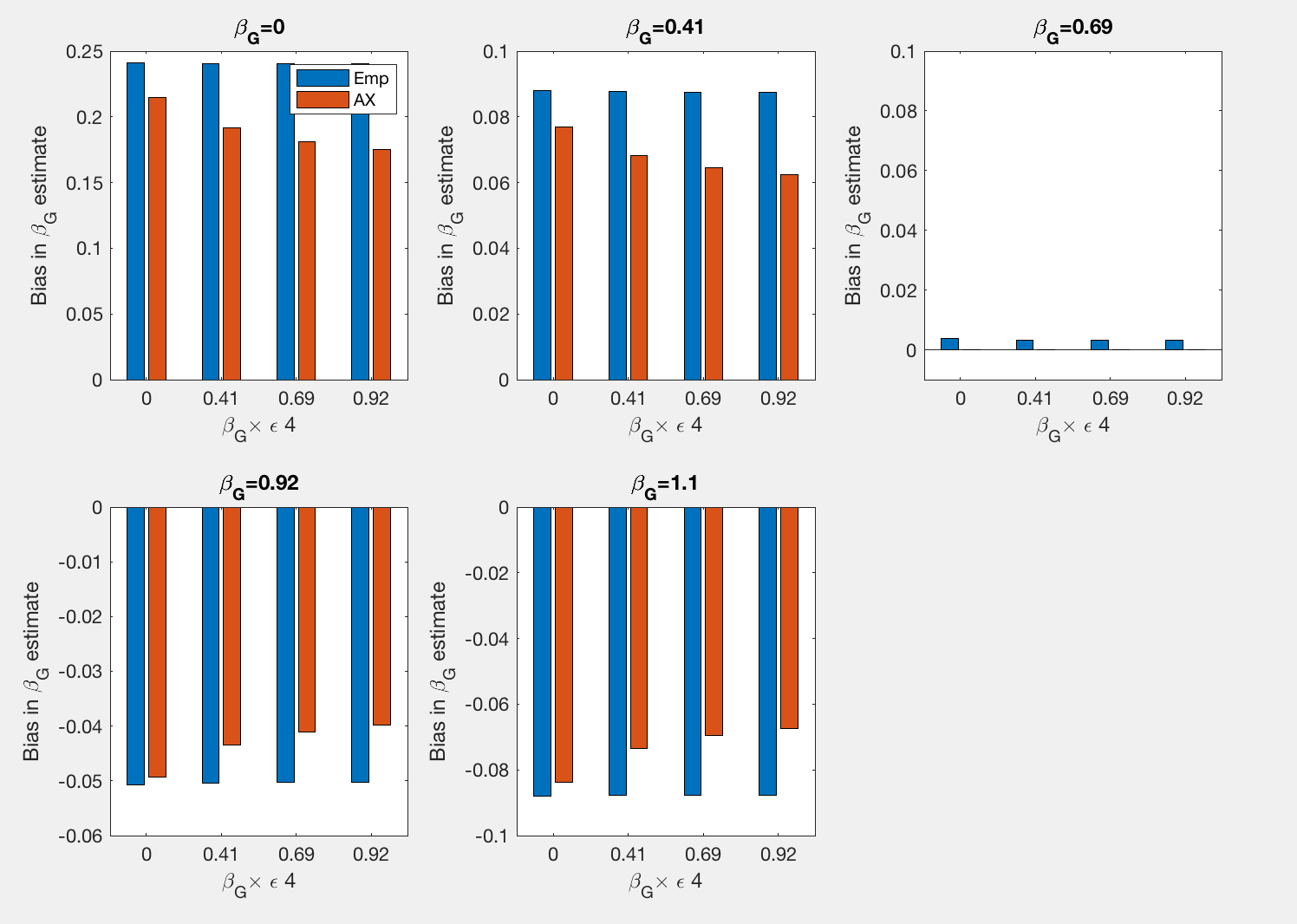

**Supplementary Figure 5:** Empirical bias (Emp) and approximation (A6c) (AX) in estimates of $\beta_{A\times\epsilon4}$ across 500 datasets with 3,000 cases and 3,000 controls, when the data are simulated using model (A3)-(A4) but the parameters are estimated using model (A5). Genotype, age, sex, ApoE $\epsilon4$ status are simulated to be binary with frequencies $\theta_{G}=0.10,\theta_{A}=0.50,\theta_{S}=0.52,\theta_{\epsilon4}=0.07$**.** The coefficients are $\beta_{0}=1.5,\beta_{S}=log(0.80),\beta_{\epsilon4}=log(8),\beta_{A}=log(3),$

$\beta_{G}=log\left( 1 \right)=0,log\left( 1.5 \right)=0.41,log\left( 2 \right)=0.69,log\left( 2.5 \right)=0.92,log\left( 3 \right)=1.1,\beta_{A\times\epsilon4}=log\left( 1 \right)=0,log\left( 1.5 \right)=0.41,log\left( 2 \right)=0.69,log\left( 2.5 \right)=0.92,\alpha_{0}=0.5,\alpha_{S}=log\left( 0.80 \right),\alpha_{\epsilon4}=log\left( 4 \right),\alpha_{A}=log\left( 3 \right),\alpha_{G}=log\left( 2 \right),\alpha_{A\times\epsilon4}=log\left( 2 \right).$ Values of $\beta_{A\times\epsilon4}$ are along the x-axis and values of $\beta_{G}$. Empirical estimates are in blue and the approximations are in red.

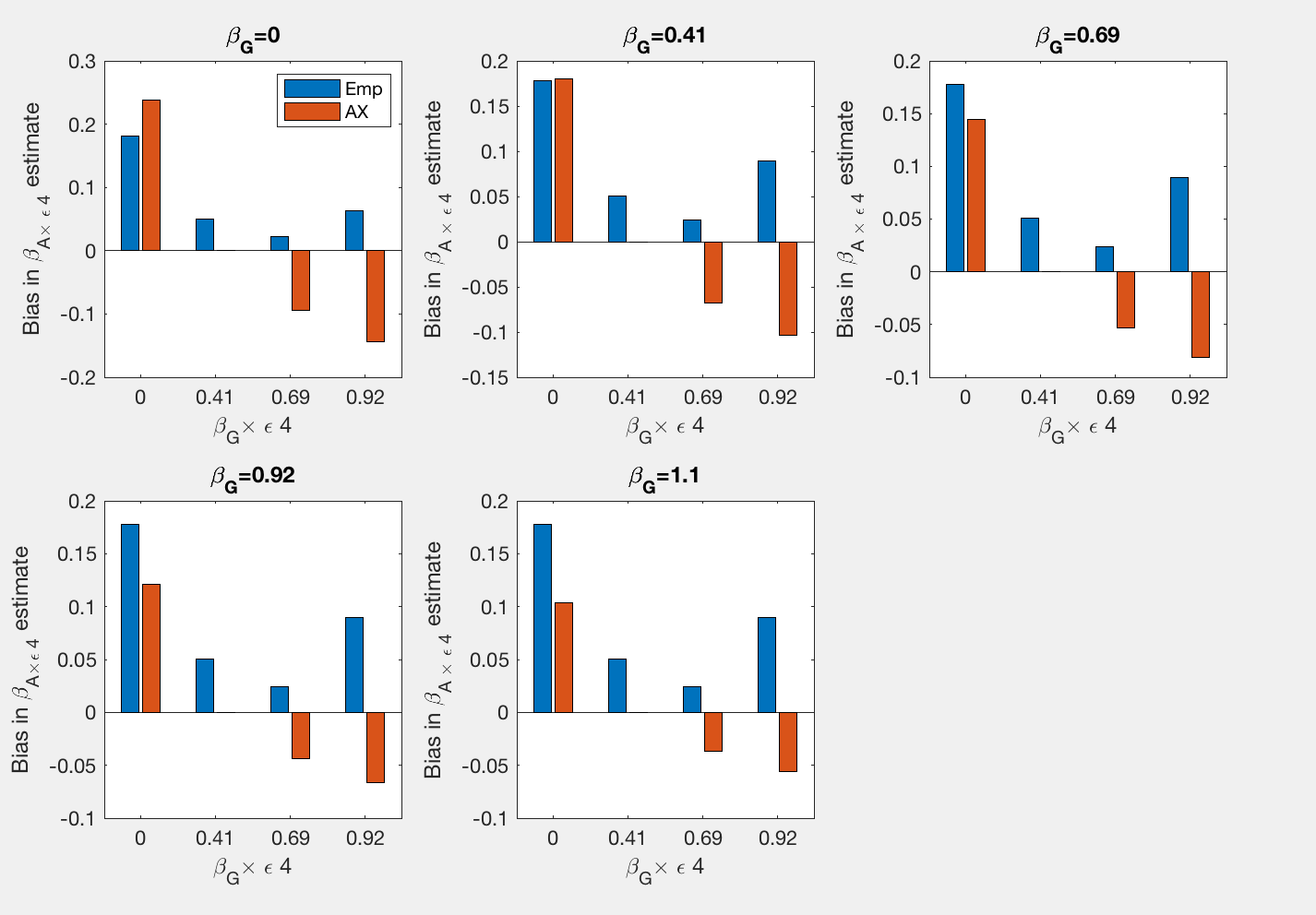

**Supplementary Figure 6:** Empirical bias (Emp) and approximation (A6c) (AX) in estimates of $\beta_{\epsilon4}$ across 500 datasets with 3,000 cases and 3,000 controls, when the data are simulated using model (A3)-(A4) but the parameters are estimated using model (A5). Genotype, age, sex, ApoE $\epsilon4$ status are simulated to be binary with frequencies $\theta_{G}=0.10,\theta_{A}=0.50,\theta_{S}=0.52,\theta_{\epsilon4}=0.07$**.** The coefficients are $\beta_{0}=1.5,\beta_{S}=log(0.80),\beta_{\epsilon4}=log(8),\beta_{A}=log(3),$

$\beta_{G}=log\left( 1 \right)=0,log\left( 1.5 \right)=0.41,log\left( 2 \right)=0.69,log\left( 2.5 \right)=0.92,log\left( 3 \right)=1.1,\beta_{A\times\epsilon4}=log\left( 1 \right)=0,log\left( 1.5 \right)=0.41,log\left( 2 \right)=0.69,log\left( 2.5 \right)=0.92,\alpha_{0}=0.5,\alpha_{S}=log\left( 0.80 \right),\alpha_{\epsilon4}=log\left( 4 \right),\alpha_{A}=log\left( 3 \right),\alpha_{G}=log\left( 2 \right),\alpha_{A\times\epsilon4}=log\left( 2 \right).$ Values of $\beta_{A\times\epsilon4}$ are along the x-axis and values of $\beta_{G}$. Empirical estimates are in blue and the approximations are in red.

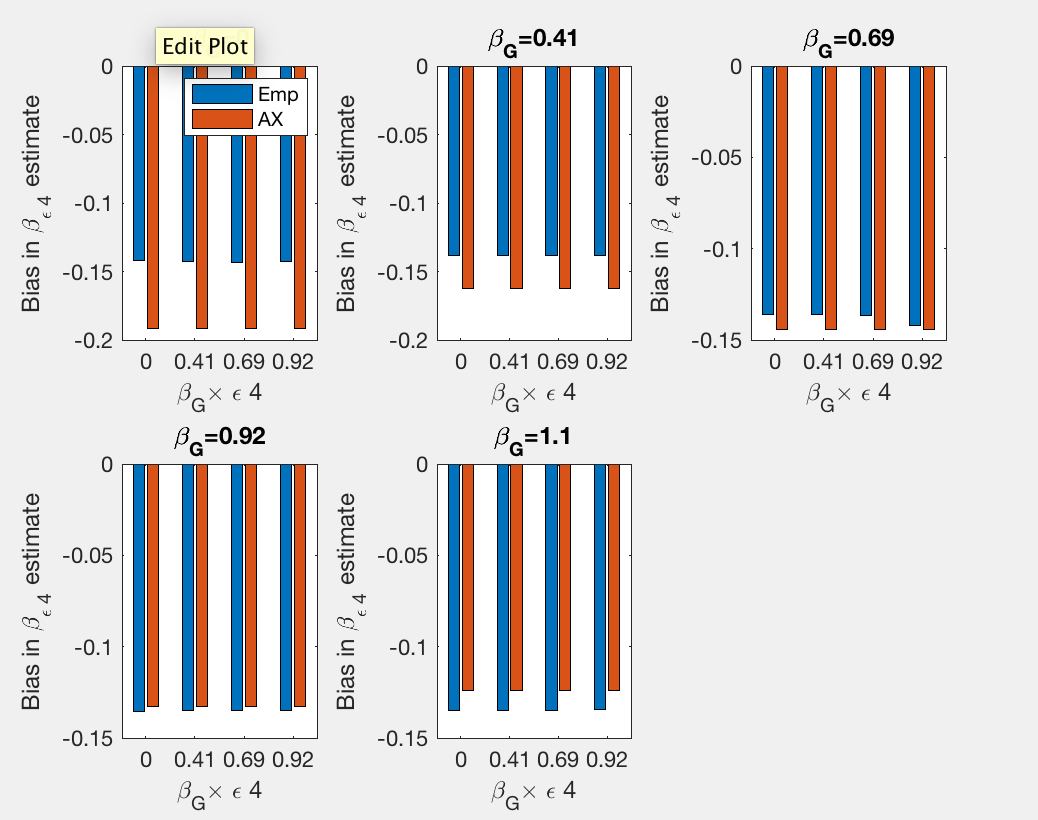

**Supplementary Figure 7:** Frequency of the disease state of interest ($D=1$) and the nuisance disease ($D=1^{*}$) when $\beta_{0}=1.5,\beta_{S}=log(0.80),\beta_{\epsilon4}=log(8),\beta_{A}=log(3),$

$\beta_{G}=log\left( 1 \right)=0,log\left( 1.5 \right)=0.41,log\left( 2 \right)=0.69,log\left( 2.5 \right)=0.92,log\left( 3 \right)=1.1,\beta_{A\times\epsilon4}=log\left( 1 \right)=0,log\left( 1.5 \right)=0.41,log\left( 2 \right)=0.69,log\left( 2.5 \right)=0.92, \alpha_{0}=0.5,\alpha_{S}=log(0.80),\alpha_{\epsilon4}=log(4),\alpha_{A}=log(3),\alpha_{G}=log(2),\alpha_{A\times\epsilon4}=log(2),$ $\theta_{G}=0.10,\theta_{A}=0.50,\theta_{S}=0.52,\theta_{\epsilon4}=0.07$**.**

Shown along the x-axis are values of $\beta_{G}$ and indicated by color are values of $\beta_{A\times\epsilon4}$.

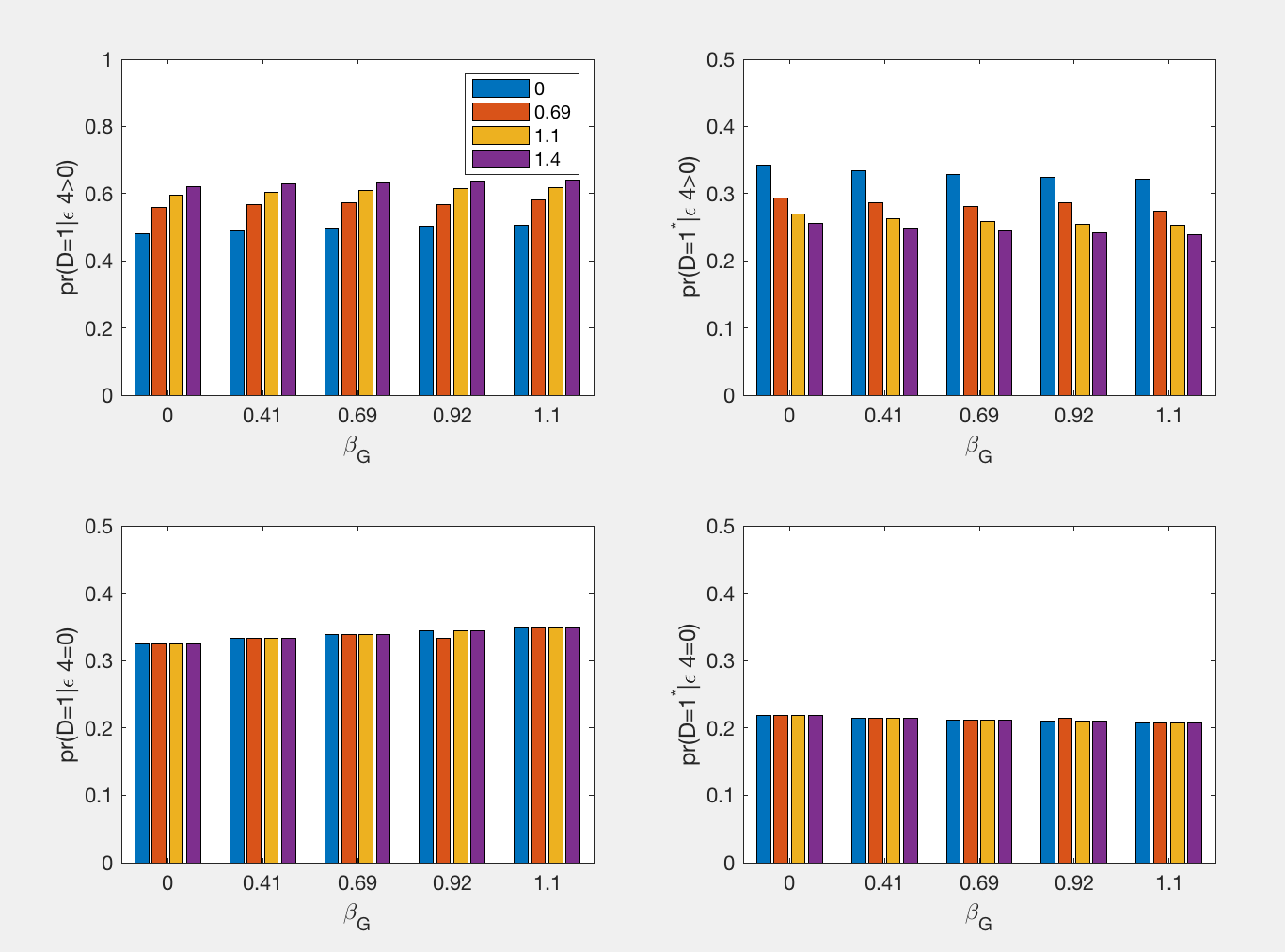
